# Single-cell and spatial transcriptomics resolve airway obliteration in bronchiolitis obliterans syndrome

**DOI:** 10.64898/2026.08.21.746071

**Authors:** Jannik Ruwisch, Hande Yilmaz, Leonard Christian, Lavinia Neubert, Lena M. Leiber, Annalea Brüggemann, Sneha Banerjee, Mark Greer, Wiebke Rackwitz, Leon Giercke, Christopher Werlein, Carina A Pawlow, Regina Engelhardt, Axelle Coppens, Matthias Ballmaier, Evgeny Chichelnitskiy, Susanne Simon, Jawad Salman, Khalil Aburahma, Ali Önder Yildirim, Janine Gote-Schniering, Jens Hohlfeld, Bard Vanaudenaerde, Danny D. Jonigk, Sabine Dettmer, Fabio Ius, Marius M Hoeper, Svenja Gaedcke, Naftali Kaminski, Yang Li, Stijn E Verleden, Jens Gottlieb, Christine Falk, Jan C. Kamp, Jonas C. Schupp

## Abstract

**Background:** Chronic lung allograft dysfunction (CLAD) is the leading cause of death beyond the first year after lung transplantation, and its most frequent phenotype is bronchiolitis obliterans syndrome (BOS), a fibrotic small-airway disease. Mechanistic work has focused on the immune compartment, yet intensified immunosuppression does not alter established disease.

**Aim:** To resolve which structural cell states populate the BOS graft and how they are spatially organized during airway obliteration.

**Methods:** We profiled explanted lungs from 33 BOS patients undergoing re-transplantation and 33 controls, combining single-nucleus RNA sequencing (14 BOS, 13 controls) with targeted spatial transcriptomics of 108 regions (27 BOS, 24 controls) and multiplex immunofluorescence validation. Single-nucleus data were integrated with a published restrictive allograft syndrome (RAS) atlas.

**Results:** Across 175,128 nuclei and 1.67 million spatially resolved cells, BOS lungs harbored a profibrotic circuit of Aberrant Basaloid cells and *CTHRC1*+ fibrotic fibroblasts previously described in fibrotic lung diseases, including RAS. Spatial mapping identified a *CXCL14*+*TNC*+ injury-associated basal cell state arising early in the obliterative cascade, identifying basal cells as their major reservoir. *CTHRC1*+ fibroblasts expanded subepithelially replacing resident peribronchial fibroblasts, alongside a peribronchial vascular shift toward systemic venous endothelium. The circuit extended beyond the airway wall to the alveolar interface, defining two convergent remodeling fronts.

**Conclusion:** BOS engages structural-cell circuits largely shared with RAS and fibrotic lung diseases, but along an airway-centered rather than parenchyma-centered axis. CLAD thus emerges as a spatial rather than cellular spectrum, defined by anatomical distribution more than cell identity. Shared structural programs may therefore be targetable across CLAD phenotypes.

## Introduction

Chronic lung allograft dysfunction (CLAD) is an irreversible decline in pulmonary function post lung transplantation that cannot be attributed to other causes. CLAD affects up to half of lung transplant (LTx) recipients within 5 years following LTx and is the leading cause of death after the first year^1^. Its most frequent phenotype, bronchiolitis obliterans syndrome (BOS), is defined by progressive airflow obstruction^2^ and histologically by fibrotic obliteration of small airways^3^. Restrictive allograft syndrome (RAS) is defined by restrictive functional changes with computed tomography-proven opacities and peripheral intraalveolar fibroelastosis (AFE). There is no effective treatment for CLAD besides re-Tx, and median survival after diagnosis remains 6–18 months in RAS and 36 months in BOS^4^.

Mechanistic work has focused almost entirely on the immune compartment: alloreactivity, donor-specific antibodies, and cytotoxic tissue-resident memory T cells^5–9^. Yet, intensifying immunosuppression does not alter the trajectory of established CLAD^4,6^ suggesting that remodeling, once initiated, is sustained by the structural compartment itself, largely independent of its immune trigger. Unresolved questions follow: Which structural cell states populate the CLAD graft and are BOS and RAS, and their diverging PFT patterns, distinct entities with discrete cellular players, or one cascade in different anatomical compartments? Phenotype coexistence and conversion argue for a shared substrate, divergent histology and survival for separate biology^7^.

Here, we analyzed explanted BOS and RAS lungs by single-nucleus RNA sequencing, focused on the structural compartment, identifying aberrant epithelial, ectopic endothelial, and fibroblast populations in BOS, and resolved their spatial distribution by spatial transcriptomics and multiplex immunofluorescence.

## Materials And Methods

Materials and methods are described only briefly here; detailed information can be found in the supplement.

### Ethics approval

This study was approved by the local Ethics Committee of Hannover Medical School (IRB # 10141_BO_K_2022).

### Sample selection

We analyzed tissue specimens of a cohort spanning 33 BOS patients undergoing re-LTx and 33 controls (CTRL; **Table 1**). For spatial analysis, we grouped s=108 samples according to the histological sequence observed in micro-CT-guided serial sections of the obliterating BOS airway, from an initial "*Unaffected*" stage through "*Inflamed*" and "*Obliterating*" to full "*Obliteration*" (**Fig.1C**), with additional samples showing the hallmarks of *AFE* remodeling (**Fig.1I-J**)^10,11^. This sample annotation is referred to as “lesion sub-class” throughout the manuscript.

**Table 1.** Detailed BOS and CTRL statistics. Statistical tests were conducted as indicated with *=*Fisher’s Exact* test for categorical and ^#^=*Wilcoxon rank-sum* test for continuous variables. HRCT patterns were evaluated and segmented by a board-certified radiologist, while histopathology patterns were reviewed by board certified pathologists. Predicted pulmonary function values were computed using Global Lung Initiative (GLI) reference equations^41^. For transplant recipients, smoking status refers to the exposure history of the graft rather than the recipient. Never-smoking recipients whose donor had a documented smoking history were therefore classified as ex-smokers.

|  | BOS (N=33) | CTRL (N=33) | Total (N=66) | P value |
| --- | --- | --- | --- | --- |
| **Sex** |  |  |  | 0.46* |
| Male | 15 (45.5%) | 19 (57.6%) | 34 (51.5%) |  |
| **Smoking status** |  |  |  | < 0.001* |
| Yes | 0 (0.0%) | 16 (48.5%) | 16 (24.2%) |  |
| Former | 13 (39.4%) | 2 (6.1%) | 15 (22.7%) |  |
| No | 20 (60.6%) | 10 (30.3%) | 30 (45.5%) |  |
| Unknown | 0 (0.0%) | 5 (15.2%) | 5 (7.6%) |  |
| **Xenium analysis** |  |  |  |  |
| True | 27 (81.8%) | 24 (72.7%) | 51 (77.3%) |  |
| **snRNAseq analysis** |  |  |  |  |
| True | 14 (42.4%) | 13 (39.4%) | 27 (40.9%) |  |
| **Tissue** |  |  |  |  |
| Explant | 33 (100.0%) | 0 (0.0%) | 33 (50.0%) |  |
| Downsizing lung | 0 (0.0%) | 25 (75.8%) | 25 (37.9%) |  |
| Peripheral tumor resection | 0 (0.0%) | 8 (24.2%) | 8 (12.1%) |  |
| **Tx modality** |  |  |  |  |
| DLTx | 33 (100.0%) | NA | 33 (50.0%) |  |
| **Clinical CLAD phenotype** |  |  |  |  |
| BOS | 25 (75.8%) | NA | 25 (37.9%) |  |
| mixed | 5 (15.2%) | NA | 5 (7.6%) |  |
| RAS | 2 (6.1%) | NA | 2 (3.0%) |  |
| Undefined | 1 (3.0%) | NA | 1 (1.5%) |  |
| CTRL | 0 (0.0%) | 33 (100.0%) | 33 (50.0%) |  |
| **Death or ReTx** |  |  |  |  |
| Yes | 13 (39.4%) | NA | 13 (39.4%) |  |
| **Azithromycin** |  |  |  |  |
| Yes | 32 (97.0%) | NA | 32 (48.2%) |  |
| **Extracorporeal photopheresis** |  |  |  |  |
| Yes | 25 (75.8%) | NA | 25 (37.9%) |  |
| **Donor Age / CTRL Age [y]** |  |  |  | 0.003# |
| Mean (SD) | 45.5 (13.8) | 54.4 (16.6) | 50.2 (15.9) |  |
| Median (Q1, Q3) | 46 (39.5, 53.8) | 59 (54, 66) | 54 (41, 63.5) |  |
| Range | 16 - 71 | 18 - 72 | 16 - 72 |  |
| **AI-Segmented Normal HRCT Volume [%]** |  |  |  |  |
| Mean (SD) | 93.8 (9.7) | NA | 93.8 (9.7) |  |
| Median (Q1, Q3) | 96.5 (92.8, 99.3) | NA | 96.5 (92.8, 99.3) |  |
| Range | 55 - 100 | NA | 55 - 100 |  |
| **FEV1 (predicted) [%]** |  |  |  |  |
| Mean (SD) | 20.9 (8.1) | NA | 20.9 (8.1) |  |
| Median (Q1, Q3) | 19 (16, 23) | NA | 19 (16, 23) |  |
| Range | 11 - 51 | NA | 11 - 51 |  |
| **FEV1 (relative to post transplant baseline) [%]** |  |  |  |  |
| Mean (SD) | 25.1 (13.1) | NA | 25.1 (13.1) |  |
| Median (Q1, Q3) | 23 (16.5, 28) | NA | 23 (16.5, 28) |  |
| Range | 9 - 76 | NA | 9 - 76 |  |
| **TLC (relative to post transplant baseline / predicted) [%]** |  |  |  |  |
| Mean (SD) | 107.2 (18.5) | NA | 107.2 (18.5) |  |
| Median (Q1, Q3) | 104.5 (98, 119.3) | NA | 104.5 (98, 119.3) |  |
| Range | 62 - 140 | NA | 62 - 140 |  |
| **Time to re-transplantation [y]** |  |  |  |  |
| Mean (SD) | 6.43 (3.80) | NA | 6.43 (3.80) |  |
| Median (Q1, Q3) | 5.91 (3.45, 8.67) | NA | 5.91 (3.45, 8.67) |  |
| Range | 0.55 - 16.20 | NA | 0.55 - 16.20 |  |
| **Time to Death [y]** |  |  |  |  |
| Mean (SD) | 9.76 (4.63) | NA | 9.76 (4.63) |  |
| Median (Q1, Q3) | 9.39 (6.72, 11.81) | NA | 9.39 (6.72, 11.81) |  |
| Range | 1.51 - 23.87 | NA | 1.51 - 23.87 |  |

**Figure 1.**
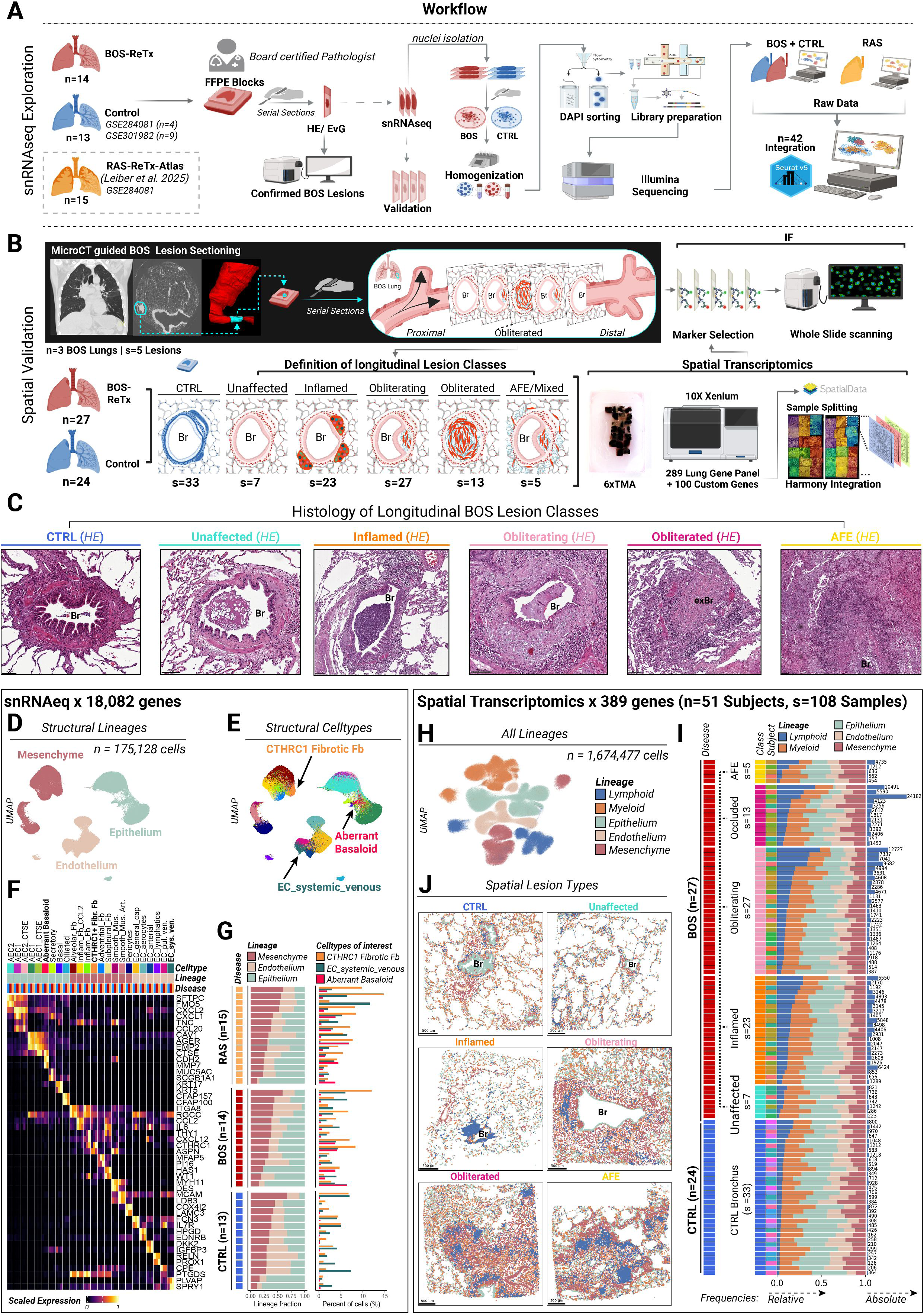
Worflow summary and dataset introduction. A) snRNAseq data was generated from 14 BOS lung re-explants. Data was integrated with snRNAseq data of n=15 RAS and 13 CTRL (GSE284081, GSE301982). B) 10X Xenium spatial transcriptomics assessing n=389 probes served as spatial validation platform. For categorization, micro-CT guided serial sectioning of s=5 lesions from s=3 BOS explants (Antwerp, Belgium), indicated distinct remodeling stages alongside the longitudinal airway-axis (lesion sub-classes), wherein s=108 samples were categorized in for subsequent comparative analysis. C) Lesion sub-classes comprised airways from CTRL patients (*CTRL*), un-remodeled BOS airways (*Unaffected*), inflamed BOS airways (*Inflamed*), obliterating airways (*Obliterating*), fully obliterated BOS airways (*Obliterated*) and BOS lesions with signs of intra-alveolar fibroelastosis in the surrounding alveoli (*AFE*). Stained with H&E. D) Uniform Manifold Approximation and Projection (UMAP) representation of structural cell lineages 175,128 nuclei, color coded by lineage. E) UMAP by cell type. Within structural cell types, we identified *CTHRC1*+ fibrotic fibroblasts, Aberrant Basaloid cells, and systemic venous endothelial cells (EC), having been previously detected across various ILDs. F) Unity scaled average expression of canonical cell type markers aggregated by disease level. G) Stacked bar plots of relative lineage frequencies per subject colored by lineage and split by cohort. Subjects were ordered after frequencies of mesenchymal cells (left panel). Bar plots denote relative cell type per-sample frequencies of established fibrotic cell types colored by cell type (right panel). H) UMAP representation of 10X Xenium data with all lineages spanning 1,674,477 cells, color coded by lineage. I) Stacked bar plots of relative lineage frequencies per sample (s=108) colored by lineage and split by lesion sub-class. Subjects were ordered after frequencies of mesenchymal cells (left panel). Bar plots denote absolute per-sample counts of lymphoid cells (right panel). J) Spatial plots of exemplary lesion sub-class colored by lineage. Scale bars denote 500 µm unless stated otherwise. Scale bars denote 500µm, unless stated otherwise. Created in BioRender. Schupp, J. (2026) https://BioRender.com/77x10o4

### snRNAseq and Spatial Transcriptomics

In brief, single nucleus RNA sequencing (snRNAseq) and spatial transcriptomics data were acquired with the Chromium Flex and Xenium platforms (both 10X Genomics), respectively. snRNAseq data were processed, integrated, clustered, and visualized with the "*Seurat*" R package and used to guide gene panel design for subsequent spatial transcriptomics. To enable phenotype-wide comparison of structural cell composition across the CLAD lung, we integrated the BOS and CTRL snRNAseq data with a recently published atlas of n=15 RAS patients^12^. For spatial transcriptomics data, the GPU-supported *scanpy* implementation *RapidSingleCell* (Python) was used due to larger data set scale. Cell type annotation was established using available cell atlases^10,12–16^ in conjunction with canonical cell type markers (**Table S1-2**). Details on data generation, processing, and analysis are provided in the Supplements.

### Validation

Molecular features were validated on protein level with multiplex immunofluorescence (IF) stains micro-CT guided, serial sectioned BOS lesions (n=3/s=5, **Fig.1B**).

### Statistics

Inter-group statistics used the *Wilcoxon rank-sum* and *Kruskal-Wallis* tests, with *post hoc* comparisons by *Wilcoxon rank-sum test* and *Benjamini-Hochberg* correction. Further details including single-cell statistics are provided in the Supplements.

## Results

### Cohort and Dataset Description

The investigated cohorts spanned n=33 BOS patients and n=33 control patients, the latter consisting of lung tissues obtained during surgical size-reduction of oversized donor lungs (n=25) and peripheral tumor resections (n=8, **Table 1**). BOS and CTRL groups were well balanced for sex (P=0.46), while BOS patients were significantly younger (P=0.003). Pulmonary function in the BOS cohort exhibited the expected obstructive pattern (median FEV1 23% of baseline, TLC 104.5%; **Table 1**). Radiologic features of inflammation or fibrosis on HRCT were absent in 96.5% of the segmented volumes. All patients met CLAD criteria with histological BOS; clinically, BOS predominated (n=25, 75.8%), followed by mixed phenotype (n=5, 15.2%) and two patients with RAS clinical criteria despite histological BOS. Median time to re-LTx was 5.9 years.

For snRNAseq-based exploratory unbiased disease profiling, we analyzed n=14 BOS, n=15 RAS, and n=13 CTRL samples, yielding a structural cell atlas of 175,128 curated nuclei (**Fig.1A**). To focus on structural remodeling, immune nuclei were excluded. snRNAseq-informed spatial transcriptomics profiling was performed on n=27 BOS and n=24 CTRL specimens using a custom-designed gene panel (Xenium, n=389 genes), applied on s=108 specimen, yielding a dataset spanning 1,674,477 cells (**Fig. 1H-I**). Per-disease statistics for lineages and cell types are provided in the Supplementary Materials, with selected findings highlighted below. Data are available on GEO (GSEXXXX) and through an interactive web portal (https://breath.mh-hannover.de/BOSatlas.html).

snRNAseq-guided lineage analysis depicted only minor variations across diseases (**Fig.1D-G**), while on lesion-level spatial analysis indicated significant frequency variations of lymphoid (Padj._KW_<0.001), endothelial (Padj._KW_=0.003) and epithelial (Padj._KW_=0.004) lineages (**Fig.1I**, **Table S3-8**). Lymphoid cells were most abundant in *Inflamed* BOS lesions, supporting the histological categorization at the cellular level (**Table S3**, **Fig.1J**). Across all structural lineages of the snRNAseq dataset (**Fig.1D**, **Table S1**), we surprisingly identified a profibrotic cell circuit in BOS previously described in fibrotic parenchymal lung diseases such as idiopathic pulmonary fibrosis (IPF)^15^, pleuroparenchymal fibroelastosis (PPFE),^10,15,17^ and RAS^12^ (**Fig.1E-G**, **Table S2**). In the following, we dissect each component of this circuit in further detail.

### Aberrant Basaloid cells emerge in the obliterating airway and line the remodeled bronchovascular bundle

Across 62,935 curated epithelial nuclei, we resolved four injury-associated states: Aberrant Basaloid cells (*MMP7*, *CDH2*, *KRT17*, *EPHB2*)^15^, *CTSE*+ inflamed AEC1 and AEC2^10,12,17^, and *CTSE*^low^ early inflamed AEC2 (*CXCL2*, *CSF3*, *CXCL1*, **Fig.2A**, **Table S2**). Of these, Aberrant Basaloid cells (Padj._KW_=0.001) were significantly enriched in both RAS and BOS lungs, while AEC1_CTSE+ exhibited numerical increase (Padj._KW_=0.074, **Fig.2B, Table S9-10**). Aberrant Basaloid (*KRT17, CTSE*, *CDH2*, *MMP7*) cells expressed their typical marker portfolio across BOS and RAS (**Fig.2C, Table S2, Suppl.Results**), with differential pseudobulk analysis confirming a conserved expression signature across both diseases (**Table S11**).

**Figure 2.**
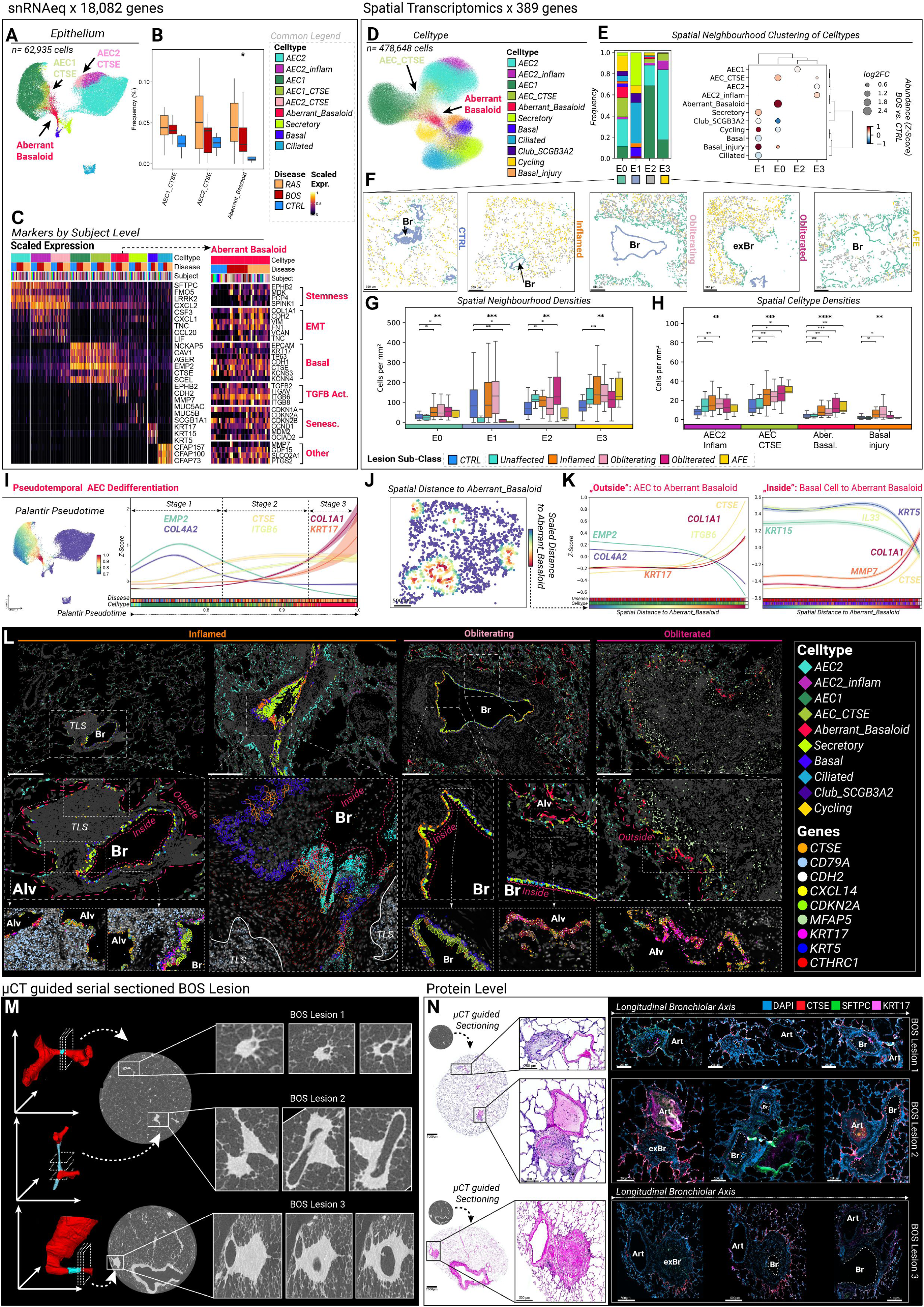
Epithelial plasticity of BOS. A) Uniform Manifold Approximation and Projection (UMAP) representation of snRNAseq data of the epithelial cell lineage 63,935 nuclei, color coded by cell type. B) Boxplots display the relative distribution of each epithelial cell type relative to the total epithelial lineage compartment, stratified by disease-related cohort. Whiskers indicate 1.5 times the interquartile range (IQR). Cohorts were compared by *Kruskal-Wallis* test, followed by *post-hoc* Wilcoxon rank sum tests, each with *Benjamini-Hochberg* correction for multiple testing. All pairwise combinations were tested, but only significant *post-hoc* comparisons versus control are displayed. Detailed results are provided in **Table S9-10**. C) Unity scaled average expression of canonical markers per subject. Subjects were grouped per disease. Each subject was assigned to a unique color. Extended differentially expressed genes of Aberrant Basaloid cells are provided as well and manually grouped after functional classes. D) Uniform Manifold Approximation and Projection (UMAP) representation of 10X Xenium data of the epithelial cell lineage spanning 478,648 curated cells, color coded by cell type. E) Stacked bar plots denote relative cell type frequency per CellCharter-related intra-epithelial spatial niche (E0-E3). The dot plot indicates enrichment of cell type per niche together with relative abundance shift comparing BOS vs. CTRL samples. F) Spatial plots of representative lesion sub-classes are colored by epithelial niche. G) Spatial densities of niche-related epithelial cells and H) cell type per computed samples area depicted as box plots per niche stratified by lesion sub-class. Whiskers indicate 1.5 times the interquartile range (IQR). Lesion sub-classes were compared by *Kruskal-Wallis* test, followed by *post-hoc* Wilcoxon rank sum tests, each with *Benjamini-Hochberg* correction for multiple testing. All pairwise combinations were tested, but only significant post-hoc comparisons versus control are displayed. Detailed results are provided in **Table S12-13**. I) Palantir pseudotime analysis analyzed gene expression trends within epithelial cells towards the Aberrant Basaloid cell state. Trends of selected significantly correlating genes are shown. Bottom bars denote cells ordered by pseudotime labeled by disease and cell type. J) Representative spatial of BOS-lesion related epithelial cells colored by scaled distance to the nearest Aberrant Basaloid cell. K) Spatially resolved expression trajectories of selected epithelial hallmark genes by scaled distance to the nearest Aberrant Basaloid cell within AEC and AEC_CTSE (left) and basal and injury-associated basal cell (right). Bottom bars denote cell type, disease label and scaled distance. L) Spatial plots of selected epithelial cell types (rhombus) and transcripts (dots) are superimposed on DAPI staining. Bronchial lumen (Br), alveolar compartment (Alv), “*In-”* and “*Outside*” sites of injury and early tertiary lymphoid structures (TLS) are annotated. M) Three-dimensional reconstructions and micro-CT images of three exemplary BOS lesions with depiction of virtual cutting planes matching serial sectioned FFPE-slides N) Serial hematoxylin-eosin and immunofluorescence (IF) staining along the longitudinal axis. Targeted sectioning was guided by micro-CT imaging of the respective tissue specimens. Bronchioles (Br), arteries (Art) and obliterated airways (exBr) are annotated Scale bars denote 500µm, unless stated otherwise.

Spatial transcriptomics confirmed the presence of *CTSE*+ alveolar epithelial cells and Aberrant Basaloid cells (**Fig.2D**, **Fig.S2A-C**, **Fig.S3, Table S12-15**). Spatially resolved Xenium analysis uncovered a BOS-associated state of *CXCL14*+ and *TNC*+ basal cells missed by snRNAseq, which we termed injury-associated basal cells (**Fig.2D**, **Table S6**). These cells were most abundant in the *Inflamed* and *Obliterating* lesions but rarely detected in *Unaffected* BOS airways or in fully *Obliterated* lesions (**Fig.2H**), consistent with an early appearance of this state during airway remodeling. Spatially, *CXCL14*+ injury-associated basal cells localized "*Inside*" to the basal epithelial layer in contact with abundant sub-epithelial *CTHRC1*+ fibrotic fibroblasts, whereas Aberrant Basaloid cells occasionally occupied the more luminal layers, coinciding with peak *MMP7* expression (**Fig.2L**).

Incorporating 2D spatial information into a cell-based niche annotation (intercellular proximity analysis and algorithmic *CellCharter*^18^ clustering), we resolved four stable epithelial niches (E0-E3, **Fig.2E**). Two of these emerged as the principal sites of epithelial dedifferentiation: the luminal bronchial epithelium (E1, "*Inside*") and the outer rim of peribronchovascular connective tissue (E0, "*Outside*"), both enriched for injury-associated basal cells and Aberrant Basaloid cells (**Fig.2E-F**). In contrast, the AEC1-(E2) and AEC2-centered (E3) niches showed only minor shifts (**Suppl.Results**). “*Outside*”, Aberrant Basaloid cells lined peribronchial connective tissue individually or as a monolayer near AEC_CTSE islets (*Obliterating*, *Obliterated* lesions) and co-localized with the B cell marker *CD79A* in some *Inflamed* lesions (light blue, **Fig.2L**), suggesting an immuno-fibrotic crosstalk explored below.

These observations were reinforced by spatial cell density analysis (per mm², **Fig.2G-H**): the "*Outside*" E0 niche showed increased density across remodeled BOS lesions relative to *CTRL* and *Unaffected* (Padj._KW_=0.004, **Fig.2G**), and both AEC_CTSE (Padj._KW_<0.001) and Aberrant Basaloid cells (Padj._KW_<0.001) increased with progressive obliteration (**Fig.2H**, **Table S12-13**). Consistently, IF staining of µCT-guided serial sections confirmed CTSE+KRT17+SFTPC+ cells emerging in the "*Outside*" compartment across all obliteration stages (s=3 lesions, **Fig.2M-N**, single channels **Fig.S4**).

Across the longitudinal lesional spectrum, Aberrant Basaloid cells reached their highest intraepithelial frequencies in fully *Obliterated* (3.9% vs. CTRL, Padj._Wilcox_*=*0.001) and *AFE* lesions (4.8% vs. *Unaffected*, Padj._Wilcox_*=*0.003) concomitant with a loss in basal (Padj._KW_=0.016) and ciliated cells (Padj._KW_=0.001, **Fig. S2**, **Table sS14-15**). This is in line with the prior snRNAseq based frequency analysis (**Fig.2B**). As these AFE lesions still derive from histologically classified BOS lungs, this suggests that AFE lesions within a *bona fide* BOS lung may represent a temporally more advanced, remodeled disease stage.

Given the occurrence of Aberrant Basaloid cells in distinct niches, we identified hallmark genes that delineate epithelial cell states along a Palantir-based pseudotemporal trajectory culminating in the Aberrant Basaloid state (**Fig.2I**) and associated their expression dynamics to a spatial proximity-to-basaloid axis (**Fig.2J-K**). This resolved epithelial dedifferentiation into three stages: 1) a homeostatic AEC1 program (*EMP2*, *COL4A2*), 2) upregulation of the damage-response marker *CTSE* together with the TGF-β-activating integrin *ITGB6*, and 3) a coordinated rise in basaloid (*KRT17*) and EMT (*COL1A1*) markers coinciding with Aberrant Basaloid annotation. Notably, this program was recapitulated spatially, with AEC1 and AEC_CTSE gene-set expressions tracking the distance to the nearest Aberrant Basaloid cell (**Fig.2K**). Likewise, injury-associated basal cells induced Aberrant Basaloid genes (*CTSE*, *COL1A1*, *CDH2*) in closer proximity to Aberrant Basaloid cells.

Altogether, epithelial transcriptional profiling delineates two cross-sectional sites of injury within the BOS airway: 1) an "*Outside*" site, where epithelial injury predominates in AEC1 and AEC2, likely giving rise to Aberrant Basaloid cells; and 2) an "*Inside*" site, where injury programs may drive basal cells to dedifferentiate into Aberrant Basaloid cells. This establishes Aberrant Basaloid cells as a conserved transcriptional state shared across parenchyma-centered (RAS, ILDs) and airway-centered (BOS) fibrotic remodeling.

### Emergence of *CTHRC1*+ Fibrotic Fibroblasts and depletion of peribronchial fibroblasts are key features of airway obliteration

snRNAseq profiling of the mesenchymal lineage (n=62,340 cells) depicted significant loss of alveolar fibroblasts (*LIMCH1*, *RGCC,* Padj._KW_=0.014) and emergence of *CTHRC1*+ fibrotic fibroblasts (*CTHRC1*, *WNT5A,* Padj._KW_=0.011)^19^, both in RAS and BOS lungs (**Fig.3A-B, Fig.S2D-F, Table S6,S16-17**). In addition, we observed the emergence of two - a CCL2+ (*CCL2*, *IL6, CXCL1*) and a CCL2-(*THY1*, *CXCL12, LUM*) - inflammatory fibroblast states^19,20^ with the latter progressively increasing along the CTRL-BOS-RAS spectrum (median 5.4% vs. 8.8% vs. 12.7%, Padj._KW_=0.007), reaching *post-hoc* significance specifically in RAS versus CTRL (Padj._Wilcox_*<*0.001) and versus BOS (Padj._Wilcox_*=*0.037, **Fig.3B**, **Table S16-17**).

**Figure 3.**
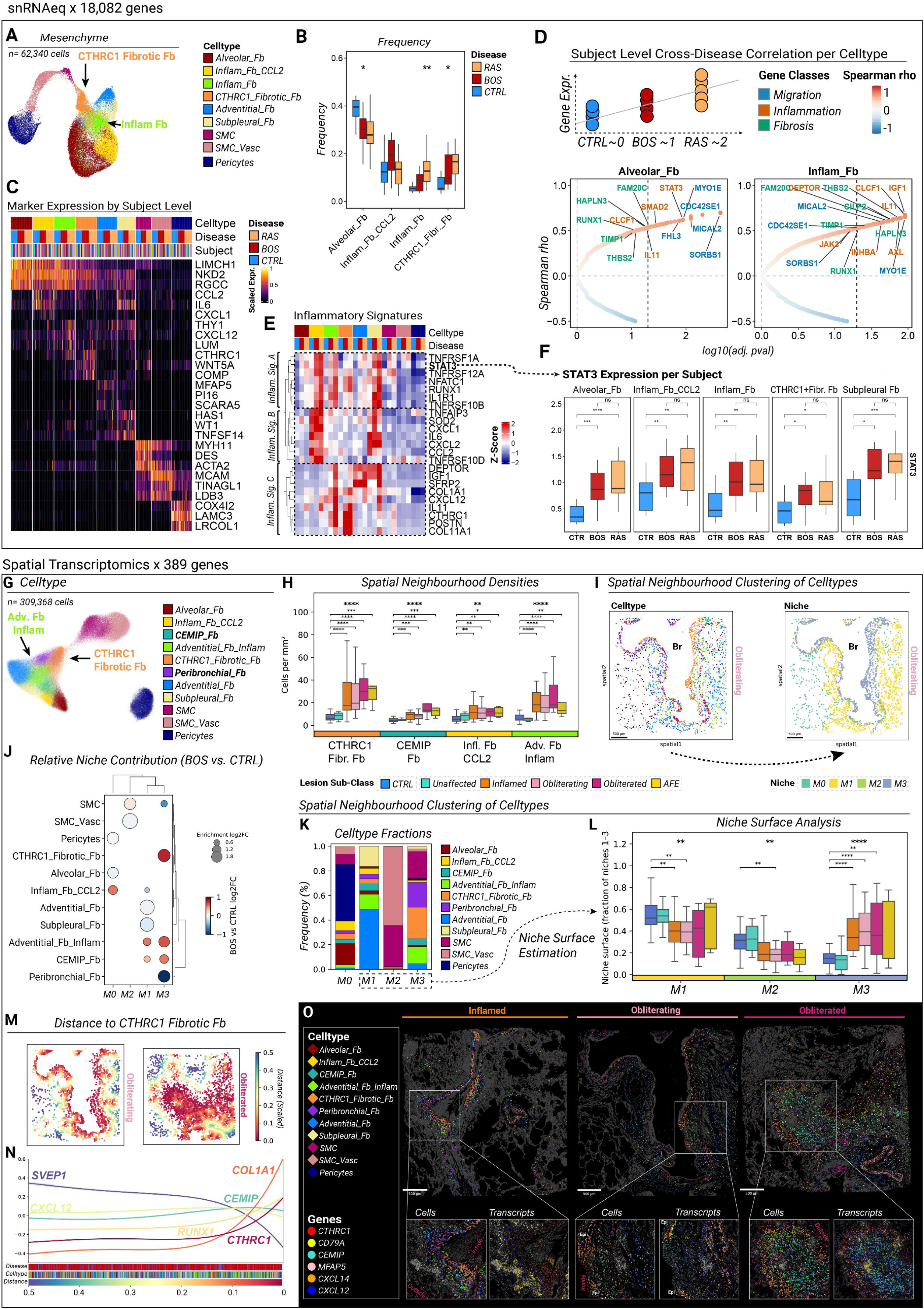
Mesenchymal plasticity of BOS. A) Uniform Manifold Approximation and Projection (UMAP) representation of snRNAseq data of the mesenchymal cell lineage 62,340 nuclei, color coded by cell type. B) Boxplots display the relative distribution of each mesenchymal cell type relative to the total mesenchymal lineage compartment, stratified by disease-related cohort. Whiskers indicate 1.5 times the interquartile range (IQR). Cohorts were compared by *Kruskal-Wallis* test, followed by *post-hoc* Wilcoxon rank sum tests, each with *Benjamini-Hochberg* correction for multiple testing. Detailed results are provided in **Table S16-17**. C) Unity scaled average expression of canonical markers per subject. Subjects were grouped per disease. Each subject was assigned to a unique color. D) Spearman correlation between subject-averaged gene expression and discrete disease encoding (0 = CTRL, 1 = BOS, 2 = RAS) across selected cell types. P values were adjusted within each cell type using the *Benjamini-Hochberg* procedure. Top-ranked genes of interest were labelled and grouped into three functional programs (Migration, Inflammation, Fibrosis). E) Scaled expression of selected significantly correlated genes, shown per cell type and collapsed to disease level. Clustering resolved three distinct inflammatory (inflam.) signatures (Sig.). F) Boxplots denote expression levels of *STAT3* averaged per subject. Inter-cohort comparisons were assessed by the *Wilcoxon Rank Sum test*. G) Uniform Manifold Approximation and Projection (UMAP) representation of 10X Xenium data of the mesenchymal cell lineage spanning 309,368 curated cells, color coded by cell type. H) Spatial densities of niche-related mesenchymal cells per computed samples area depicted as box plots per niche stratified by lesion sub-class. Whiskers indicate 1.5 times the interquartile range (IQR). Lesion sub-classes were compared by *Kruskal-Wallis* test, followed by *post-hoc* Wilcoxon rank sum tests, each with *Benjamini-Hochberg* correction for multiple testing. All pairwise combinations were tested, but only significant post-hoc comparisons versus control are displayed. Detailed results are provided in **Table S12-13**. I) Spatial plots illustrating representative cell type-to niche encoding with plots colored by mesenchymal cell type and niche (M0-M3). J) The dot plot indicates enrichment of cell type per niche together with relative abundance shift comparing BOS vs. CTRL samples. K) Stacked bar plots denote relative cell type frequency per *CellCharter*-related intra-mesenchymal spatial niche (M0-M3). L) Box plots showing the relative surface proportions of the three niches constituting the broncho-vascular bundle, renormalized to sum to 100% per sample. Boxes indicate the median and interquartile range (IQR); whiskers extend to 1.5xIQR, and each point represents one sample). Lesion sub-classes were compared by *Kruskal-Wallis* test, followed by *post-hoc* Wilcoxon rank sum tests, each with *Benjamini-Hochberg* correction for multiple testing. All pairwise combinations were tested, but only significant post-hoc comparisons versus control are displayed. Detailed results are provided in **Table S20**. M) Representative spatial of BOS-lesion related mesenchymal cells colored by scaled distance to the nearest *CTHRC1*+ fibrotic fibroblast. N) Spatially resolved expression trajectories of selected mesenchymal hallmark genes by scaled distance to nearest *CTHRC1*+ fibrotic fibroblast. Bottom bars denote cell type, disease label, and scaled spatial distance. O) Spatial plots of selected mesenchymal cell types (rhombus) and transcripts (dots) superimposed on DAPI staining, with former epithelial cells (Epi) being annotated (dashed line). Scale bars denote 500µm, unless stated otherwise.

To characterize inflammatory fibroblast plasticity in the CLAD lung, we performed an ordinal trend analysis across the CTRL–BOS–RAS spectrum analysis (**Fig.3C**), identifying genes with consistent monotonic regulation that mapped to "migration", "fibrosis", and "inflammation" programs. Hierarchical clustering of these genes in inflammatory fibroblasts resolved three inflammatory signatures (A–C) with distinct fibroblast-subtype specificity and disease-course dynamics (**Fig.3C**, Supp.Results). Most notably, the conserved signature A (*TNFRSF1A*, *TNFRSF12A*, *STAT3*) increased progressively from CTRL to BOS to RAS, with per-subject pseudobulk analysis confirming significant *STAT3* upregulation in both BOS and RAS compared to CTRL in various fibroblast populations (**Fig.3D,Table S11**).

Targeted spatial transcriptomics confirmed emergence of *CTHRC1*+ fibrotic (*CTHRC1*, *WNT5A*), *CCL2*+ inflammatory (*CCL2*, *ITGA8*), and *CCL2*-inflammatory adventitial fibroblasts (*CXCL12*, *SFRP2*) in BOS lesions (**Fig.3G, Fig.S2,Table S6**), and delineated two additional populations (bold labels, **Fig.3G**): 1) peribronchial fibroblasts (*LGR5*, *DIO2*)^10,19,21,22^ and 2) a population of migratory *CEMIP* (*Cell Migration-inducing hyaluronidase 1*)-expressing fibroblasts (*CEMIP*, *RGCC*). Both spatial density and proportion analyses confirmed the snRNAseq finding (**Fig.3B**): *CTHRC1*+ fibrotic fibroblasts increased along the longitudinal remodeling continuum (Padj._KW_<0.001), peaking in fully *Obliterated* and *AFE* lesions vs. *CTRL* (each Padj._Wilcox_<0.001, **Fig.3H**, **Table S12-13**), with *CCL2*-adventitial, *CEMIP*+ and *CCL2*+ inflammatory fibroblasts showing similar trends (all Padj._KW_≤0.001, **Fig.3H**, **Table S12-13**). Proportionally, *CTHRC1*+ fibrotic and *CCL2*-adventitial fibroblasts likewise increased with remodeling degree (both Padj._KW_<0.001, **Fig.S2**, **Table S18-19**), whereas *CEMIP*+ fibroblast proportions were highest in *Obliterated* lesions (median 4.6%, Padj._Wilcox_ vs. *CTRL*=0.004), while *Inflamed* and *Obliterating* lesions showed proportions comparable to CTRL (median 3.2%, 3.4%, and 3.0%, respectively) (**Fig.S2**).

*CellCharter*-based mesenchymal neighborhood clustering (**Fig.3I**) resolved four mesenchymal tissue niches (M0–M3, **Fig.3J-K**): an alveolar stromal niche (M0), the loose connective-tissue niche of the bronchovascular bundle (M1), the perivascular smooth muscle niche (M2), and a subepithelial niche directly beneath the luminal airway epithelium (M3). Across niches, BOS remodeling was accompanied by expansion of inflammatory and fibrotic fibroblast states (*CCL2*+ inflammatory, *CCL2*-inflammatory adventitial, *CTHRC1*+ fibrotic, and *CEMIP*+ fibroblasts) at the expense of homeostatic populations (**Fig.3J**, Supp.Results). The subepithelial M3 niche was of particular interest given its close proximity to *CXCL14*+ injury-associated basal and Aberrant Basaloid cells. In *CTRL* airways, it comprised primarily peribronchial fibroblasts, whereas BOS lesions showed a marked expansion of *CTHRC1*+ fibrotic fibroblasts, *CCL2*-inflammatory adventitial fibroblasts and, to a lesser extent, *CEMIP*+ fibroblasts, mirrored by a reciprocal BOS-related depletion of peribronchial fibroblasts (**Fig.3J**). Supporting its pathogenic relevance, M3 expanded spatially within the bronchovascular bundle (M1–3) across all BOS remodeling stages (Padj._KW_<0.001), reaching highest surface proportions in *Obliterating* lesions (median 39.3%, Padj._Wilcox_ vs. CTRL <0.001, **Fig.3L**, **Table S20**).

Because *CTHRC1*+ fibrotic fibroblasts constituted the largest fraction of the M3 niche, we asked which gene programs precede this fibrotic activation. For each fibroblast, we computed the spatial distance to the nearest *CTHRC1*+ fibrotic fibroblast (**Fig.3M**). Profibrotic markers (*COL1A1*, *CTHRC1*, *RUNX1*) were progressively upregulated in neighboring stromal cells along this distance axis, preceded by a slight rise in *CEMIP* and *CXCL12* (**Fig.3N**). Spatially, CTHRC1+ fibrotic fibroblasts formed subepithelial aggregates at the "*Inside*" remodeling edge in Inflamed and Obliterating lesions (**Fig.3O**, **Fig.S5**), with sparse inflammatory adventitial and peribronchial fibroblasts located more deeply. In *Obliterated* lesions, these cells were also located adjacent to the overlying injury-associated alveolar epithelial cell states of the "*Outside*" niche (**Fig.2L, 3O**), in spatial association with inflammatory adventitial and peribronchial fibroblasts, whereas *CEMIP*+ fibroblasts accumulated within the former bronchial lumen (**Fig.3O**). At the transcript level, dense *CD79A* aggregates were located adjacent to *CXCL12* and directly beneath subepithelial *CTHRC1*, consistent with an inflammatory-to-fibrotic gene expression gradient extending from the adventitia towards the subluminal stroma.

### Systemic venous vasculature is expanded in the perilesional connective tissue

Untargeted snRNAseq analysis of the endothelial lineage revealed a relative increase in systemic venous EC (*PLVAP, COL15A1*) in BOS (median 15.9%) and RAS (14.5%) compared to CTRL (8.1%) that did not reach statistical significance (Padj.Wilcox=0.374 for each comparison), concomitant with a numeric depletion of aerocytes (*HPGD*, *EDNRB*, **Fig.4A-C, Table sS21-22**). Lymphatic EC (*CCL21*, *PROX1*) showed no fluctuations in frequency across cohorts (**Fig.4B**, **Table S21-22**). Targeted lesional spatial transcriptomics confirmed the presence of all major endothelial cell types across the 261,657 curated EC (**Fig.4D**). However, densities of systemic venous EC per samples were drastically increased in remodeled BOS lesions (Padj._KW_<0.001, **Table S12-13**, **Fig.4E**). Density based quantities were further supported by proportional cell frequency analysis demonstrating significantly increased cell proportions in *Inflamed* and Obliterating BOS lesions (median 19% and 20.6%, respectively, Padj._Wilcox_*=*0.049 and 0.039, respectively) compared to CTRL (median 14.2%, **Table S23-24**).

**Figure 4.**
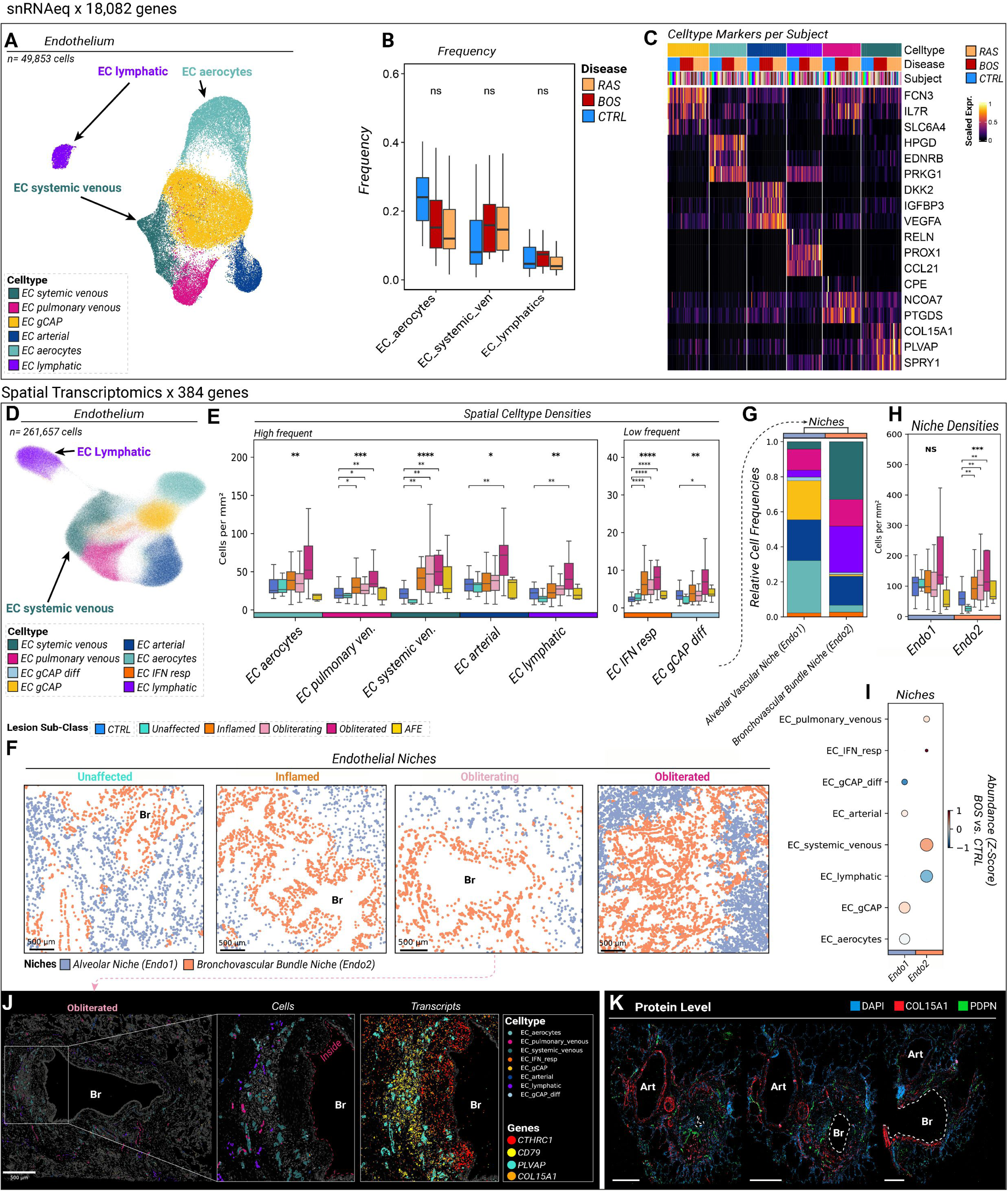
Endothelial plasticity of BOS. A) Uniform Manifold Approximation and Projection (UMAP) representation of snRNAseq data of the endothelial cell lineage 49,853 nuclei, color coded by cell type. B) Boxplots display the relative distribution of each selected endothelial cell types relative to the total endothelial lineage compartment, stratified by disease. Whiskers indicate 1.5 times the interquartile range (IQR). Cohorts were compared by *Kruskal-Wallis* test, followed by *post-hoc* Wilcoxon rank sum tests, each with *Benjamini-Hochberg* correction for multiple testing. Detailed results are provided in **Table S21-22**. C) Unity scaled average expression of canonical markers per subject. Subjects were grouped per disease. Each subject was assigned to a unique color. D) Uniform Manifold Approximation and Projection (UMAP) representation of 10X Xenium data of the endothelial cell lineage spanning 261,657 curated cells, color coded by cell type. E) Spatial densities of endothelial cell types per computed samples area depicted as box plots per niche stratified by lesion sub-class. Whiskers indicate 1.5 times the interquartile range (IQR). Lesion sub-classes were compared by *Kruskal-Wallis* test, followed by *post-hoc* Wilcoxon rank sum tests, each with *Benjamini-Hochberg* correction for multiple testing. All pairwise combinations were tested, but only significant post-hoc comparisons versus control are displayed. Detailed results are provided in **Table S12-13**. F) Spatial plots illustrate intra-endothelial niche encodings with plots (Endo1 and Endo2). G) Stacked bar plots denote relative cell type frequency per *CellCharter*-related intra-endothelial spatial niche (Endo1 and Endo2). H) Spatial densities of niche-related endothelial cells per computed samples area depicted as box plots per niche stratified by lesion sub-class. Whiskers indicate 1.5 times the interquartile range (IQR). Lesion sub-classes were compared by *Kruskal-Wallis* test, followed by *post-hoc* Wilcoxon rank sum tests, each with *Benjamini-Hochberg* correction for multiple testing. All pairwise combinations were tested, but only significant *post-hoc* comparisons versus control are displayed. I) The dot plot indicates enrichment of cell type per niche together with relative abundance shift comparing BOS vs. CTRL samples. J) Spatial plots of selected endothelial cell types (rhombus) and transcripts (dots) are superimposed on DAPI staining. With bronchiolar lumen annotated as (Br). K) Serial immunofluorescence (IF) staining along the longitudinal axis. Targeted sectioning was guided by micro-CT imaging of the respective tissue specimens. Bronchioles (Br) and arteries (Art) are annotated Scale bars denote 500µm, unless stated otherwise.

Spatial neighborhood analysis allowed us to separate the alveolar (Endo1) from the bronchovascular niche (Endo2, **Fig.4F**). As expected, lymphatic EC and systemic venous EC predominated in the latter niche (**Fig.4G**). Importantly, densities of Endo2-cells were significantly elevated in remodeled BOS airways (Padj._KW_<0.001), accompanied with a disproportionate gain in systemic venous over lymphatic EC in BOS samples (**Fig.4I-J, Table S23-24**). Sequential IF staining of the same obliterating BOS airway confirmed that COL15A1+ systemic vessels surrounded the airway, co-localized with aggregates of small DAPI+ nuclei suggestive of lymphocyte clusters and protruded into the connective tissue narrowing the airway lumen (**Fig.4K**).

### BOS lesions can be deconstructed into seven cellular niches

For pan-lineage decomposition of BOS lesions into cellular niches, we again applied the *CellCharter* workflow following comprehensive lymphoid and myeloid cell annotation (**Fig.S8, Table S6**), identifying seven major tissue niches (N1–N7, **Fig.5A**) at the most stable clustering resolution (k=7, Supp.Results). The *Bronchial Epithelial Niche* (N1) comprised airway epithelial cells and showed the strongest BOS-related shift in injury-associated basal cells, with induction of the injury markers *TNC* and *CXCL14* and emergence of Aberrant Basaloid-related genes (*FN1*, *CDH2*, *ITGB6*, **Fig.5B-D**). The adjacent *Subepithelial Stromal Niche* (N2) which intersects N1 and the *Peribronchial Stromal Niche* (N4), normally harbors peribronchial fibroblasts. In BOS, it became dominated by *CTHRC1*+ fibrotic fibroblasts that form focal subepithelial protrusions populated by injury-associated basal and Aberrant Basaloid cells (**Fig.5B-C**). Spatial proximity analysis revealed a significantly closer topographic relationship between N2 and lesional immune aggregates in BOS than in CTRL (**Fig.5E**), suggesting a local subepithelial immuno-fibrotic remodeling axis.

**Figure 5.**
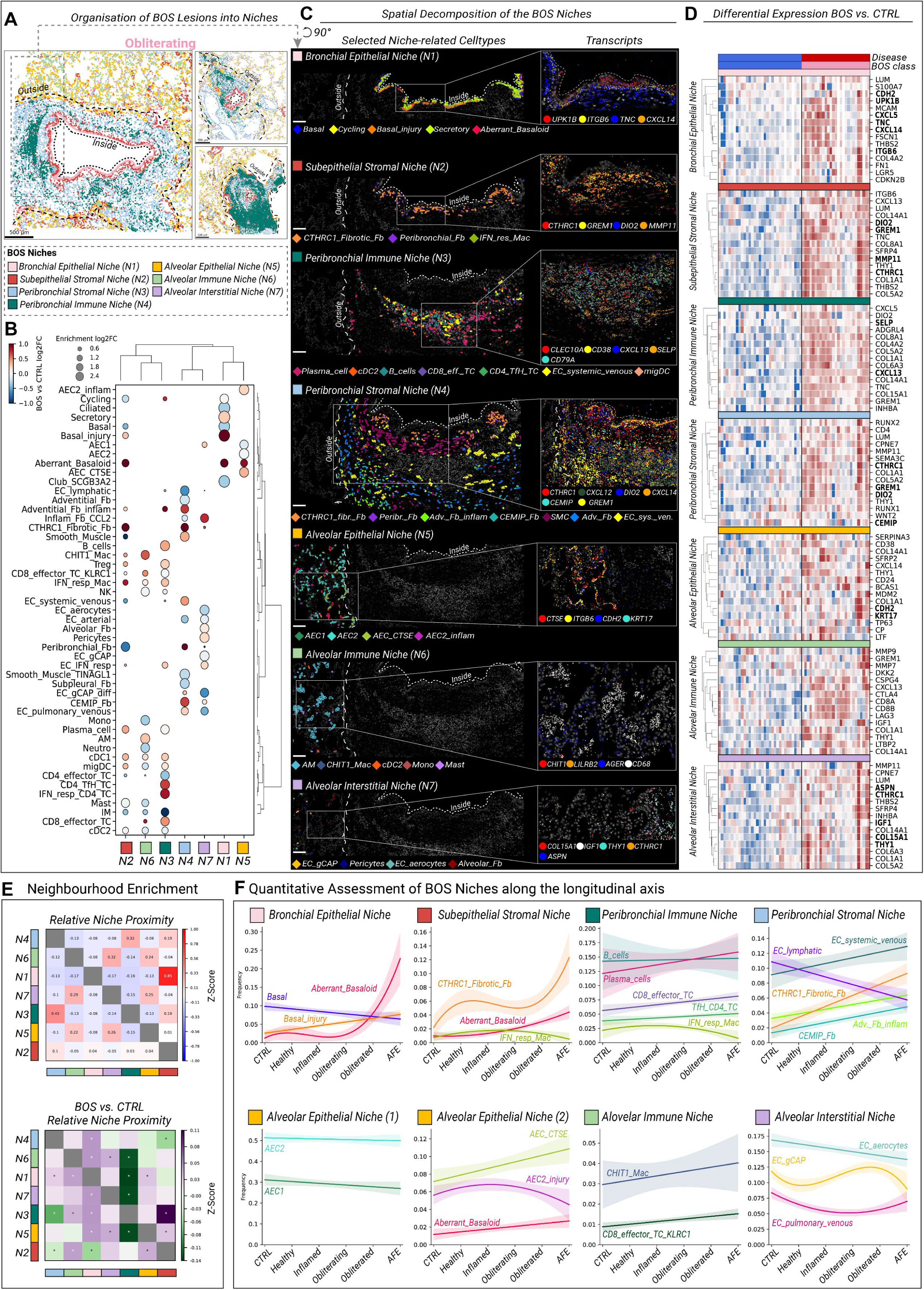
Pan-lineage niches of the BOS lesion. A) Representative spatial plots illustrate pan-lineage BOS niches (N1-N7) resolved by *CellCharter* analysis in *Obliterating* BOS lesions. B) The dotplot indicates enrichment of cell type per niche together with relative abundance shift comparing BOS vs. CTRL samples. C) Spatial Zoom-Ins demonstrating spatial localization of selected cell types (rhombus) and transcripts (dots) superimposed on DAPI staining per CellCharter-resolved niche. The “*Outside*” (coarse dashed line) and “*Inside*” (fine dashed line) remodeling sites are indicated. D) Heatmaps show niche-level pseudobulk differential expression (DE) computed in *edgeR* by contrasting each lesion sub-class against CTRL within the same niche, with *Benjamini-Hochberg*-adjusted P values. The top 15 significant DE genes (*Obliterating* vs. *CTRL*) are shown, with highlighted genes visualized in C). E) *CellCharter*-based inter-niche spatial neighborhood enrichment analysis, with proximities colored by z-score (upper panel) and differential enrichment (BOS vs. CTRL) colored by differential z-score (violet = higher adjacency in BOS; green = higher adjacency in CTRL). F) Per-niche cell-type frequencies were modeled along an ordinal severity axis (*CTRL*, *Unaffected*, *Inflamed*, *Obliterating*, *Obliterated*, *AFE*) by weighted least-squares polynomial regression, weighting samples by niche cell number and selecting the polynomial degree (linear to cubic) by lack-of-fit F-test. Fitted mean curves are shown with 95% confidence bands. Scale bars denote 100µm, unless stated otherwise.

The *Peribronchial Immune Niche* (N3) localized near N4 and formed lymphoid aggregates resembling early-stage tertiary lymphoid structures (TLS)^23–25^, particularly in *Inflamed* and *Obliterating* lesions, alongside *SELP*+ systemic venous EC as a putative recruitment site (**Fig.5B**-**C**,**E**). *The Peribronchial Stromal Niche* (N4) engulfed these TLS and showed high concentrations of inflammatory fibroblast-derived *CXCL12* (**Fig.4C**), a known driver of TLS formation^10,26,27^. Together with increased inflammatory adventitial and *CTHRC1*+ fibrotic fibroblasts and upregulation of the profibrotic transcription factors *RUNX1* and *RUNX2* and ECM-remodeling enzymes (**Fig.4B-D**), this indicates a fibrotic transformation paralleling that of the subepithelial compartment. In the alveolar compartment, the *Alveolar Epithelial Niche* (N5) surrounding the bronchovascular connective tissue was dominated by *CTSE*+ and *ITGB6*+ AECs with a strong increase in Aberrant Basaloid cells, whereas the *Alveolar Immune* (N6) and *Alveolar Interstitial* (N7) *niches* were enriched for *CHIT1*+ interstitial macrophages, alveolar macrophages, and *CCL2*+ inflammatory fibroblasts in BOS relative to CTRL (**Fig.5B-C**).

Together, these inflammation-associated cell types across niches N5–N7, alongside the inflammatory-fibrotic airway remodeling, further support the concept of two fronts of ongoing inflammatory insult in the BOS lung – an "*Inside*" lesion originating at the airway lumen and an "*Outside*" lesion in the alveolar compartment – that may converge to drive fibrotic obliteration of the BOS airway.

### Stage-resolved *Inside* and *Outside* remodeling of the BOS lesion

Exploring the longitudinal bronchial remodeling axis (treating lesion sub-classes as equidistant units of increasing severity), we observed within the "*Inside*" niches (N1-N4) a progressive emergence of an immune-fibrotic cell circuit comprising injury-associated basal cells, Aberrant Basaloid cells, inflammatory adventitial fibroblasts, *CEMIP*+ fibroblasts, *CTHRC1*+ fibrotic fibroblasts, and a TLS-associated immune cell repertoire (**Fig.5F**, upper panel). In parallel, canonical basal cells and lymphatic EC were depleted; the latter showing a reciprocal relationship with systemic venous EC already apparent in Inflamed lesions, suggesting vascular remodeling as an early event in the obliteration cascade.

In the "*Outside*" compartment, *CCL2*+ injury-associated AEC2 peaked early at the periphery of Inflamed lesions, whereas *CTSE*+ AECs increased more continuously (**Fig.5F**, lower panel), paralleled by the emergence of *CHIT1*+ interstitial macrophages and *KLRC1*+ *CD8*+ effector T-cells (*KLRC1*, *KLRD1*, *CD8A*) in the peribronchial alveolar parenchyma. Meanwhile, homeostatic cells of the blood-gas barrier (AEC1, AEC2, and aerocytes) were depleted across the remodeling continuum.

### BOS lesions harbor intertwined inflammatory and fibrotic cell circuits

Next, we used the spatially resolved cellular architecture to infer co-localizing networks around hallmark cell types of BOS-related remodeling. Aberrant Basaloid cells emerged at two lesional sites (**Fig.6A-B**), showing the closest interepithelial proximity to injury-associated basal cells (“*Inside”* the airway) and *CTSE*+ AECs (“*Outside*” the airway), as well as to *SCGB3A2*+ club cells, a relationship also evident across CTRL lesions. In line with precision-cut lung slice perturbation studies^17^, this supports that Aberrant Basaloid cells arise from three distinct reservoirs: injured *CTSE*+ AECs (paralleling epithelial dedifferentiation in RAS^12^ and IPF^15,17^), a common *SCGB3A2*+ club cell reservoir and injured *CXCL14*- and *TNC*-expressing basal cells.

**Figure 6.**
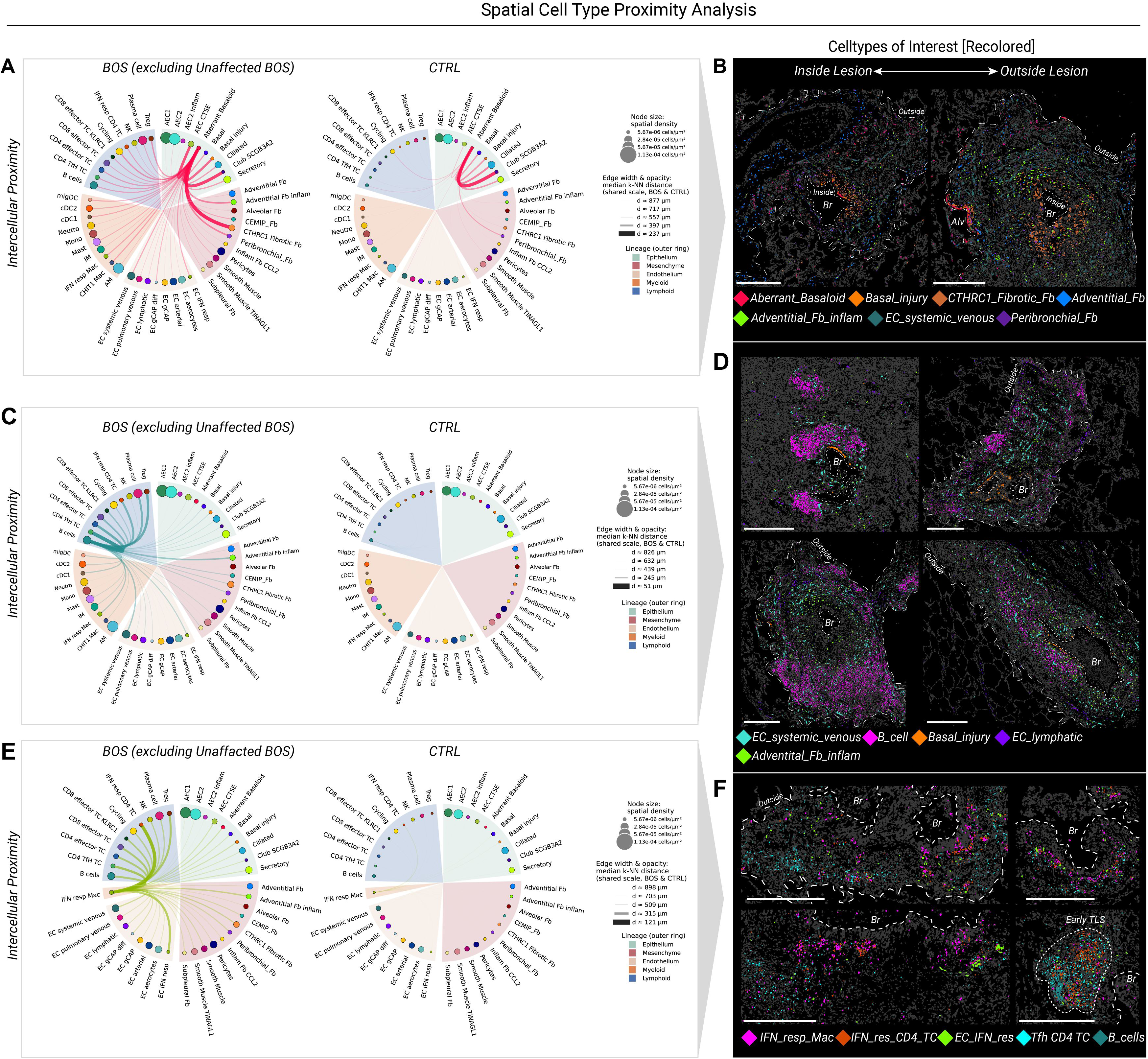
Spatially resolved BOS-related cell circuits. A) Lineage-ordered circos plot showing co-localization network centered around Aberrant Basaloid cells inferred from spatial proximity (mean Euclidean distance to the ten nearest target cells), summarized as the median per cell type and split by disease group, excluding *Unaffected* BOS samples. Node size encodes spatial density, and edge width and opacity encode median nearest-neighbor distance. B) Spatial plots demonstrate spatial localization of selected cell types co-localizing with Aberrant Basaloid cells (rhombus). The “*Outside*” (coarse dashed line) and “*Inside*” (fine dashed line) remodeling sites and bronchiolar lumen (Br) are highlighted. Cell types were partially re-colored for improved contrast. C) Lineage-ordered circos plot showing co-localization network centered around B cells analogously to A). D) Spatial plots demonstrate spatial localization of selected TLS-forming cell types, co-localizing with B cells (rhombus). The “*Outside*” (coarse dashed line) and “*Inside*” (fine dashed line) remodeling sites, early forming tertiary lymphoid structures (TLS) and bronchiolar lumen (Br) are highlighted. Cell types were partially re-colored for improved contrast. E) Lineage-ordered circos plot showing the co-localization network centered around IFN-gamma response macrophages analogously to A). F) Spatial plots demonstrate spatial localization of IFN-response cell circuits (rhombus). The “*Outside*” (coarse dashed line) and “*Inside*” (fine dashed line) remodeling sites and bronchiolar lumen (Br) are highlighted. Cell types were partially re-colored for improved contrast. Scale bars denote 500µm, unless stated otherwise.

Among the B cell-related circuits, B cells lay closest to *CXCL13*+ T follicular helper like (TfH-like) cells, supporting *CXCL13*-mediated TLS formation in BOS-related airway obliteration, and to *PLVAP*+ *COL15A1*+ systemic venous EC (**Fig.6C-D**). Given PLVAP’s role in endothelial fenestrae formation and induced *SELP* expression, this vascular cell type may serve as a primary recruiting site for local immune cells, potentially taking on the role of high endothelial venules in TLS formation. Within the mesenchymal lineage, B cells located closest to inflammatory adventitial fibroblasts expressing *CXCL12*. Consistent with this proximity, ligand-receptor analysis revealed co-localization of *CD79A* and the ECM component *FN1*, linking *CD79*+ B-cells to ECM-secreting fibroblasts as an immune-mesenchymal crosstalk (**Fig.S12**).

Finally, we identified a cellular interferon (IFN)-response hub composed mainly of IFN-activated ECs (**Fig.4D**), IFN-response macrophages and IFN-response CD4+ T cells (each with *CXCL10*, *CXCL9*, **Fig.6E,S8**), forming aggregates near early TLS-forming cells (B cells, CD4+ TfH-like T cells, **Fig.6F**) within the *Peribronchial Immune Niche* and occasionally in the *Subepithelial Stromal Niche*. IFN-response macrophage frequencies peaked in *Inflamed* and *Obliterating* lesions, suggesting IFN signaling as an early event preceding fibroblast-driven airway obliteration (**Fig.5F**).

### Unsupervised sample clustering resolves BOS lesions into four metaclusters

To assess how much of the compositional information resolved by spatial transcriptomics is already conveyed by conventional histological annotation, we re-assigned samples in a data-driven manner to four metaclusters (MC1–4) using Aitchison distances and Ward clustering (MC1–4, **Fig.7A-D**) based on computed Aitchison distances and subsequent Ward clustering. This (meta-)classification increased the explained compositional cell type variance from 0.21 (adj. 0.17) to 0.34 (adj. 0.32, **Fig.7E**) and sharpened group separation across the cellular landscape (**Fig.7F**). Meanwhile, the clinical PFT-related classification (BOS vs. Mixed vs. RAS) had only modest effects on cell compositional variance of few cell types (smooth muscle, *CEMIP*+ and inflammatory adventitial fibroblasts, **Fig.7G**, orange). The Usual Fibrotic niche (Aberrant Basaloid + *CTHRC1*+ fibrotic fibroblasts)^10,15^ showed the highest compositional variance for time to CLAD onset, with a negative correlation (**Fig.7H**), suggesting more pronounced fibrotic remodeling in patients with early CLAD onset.

**Figure 7.**
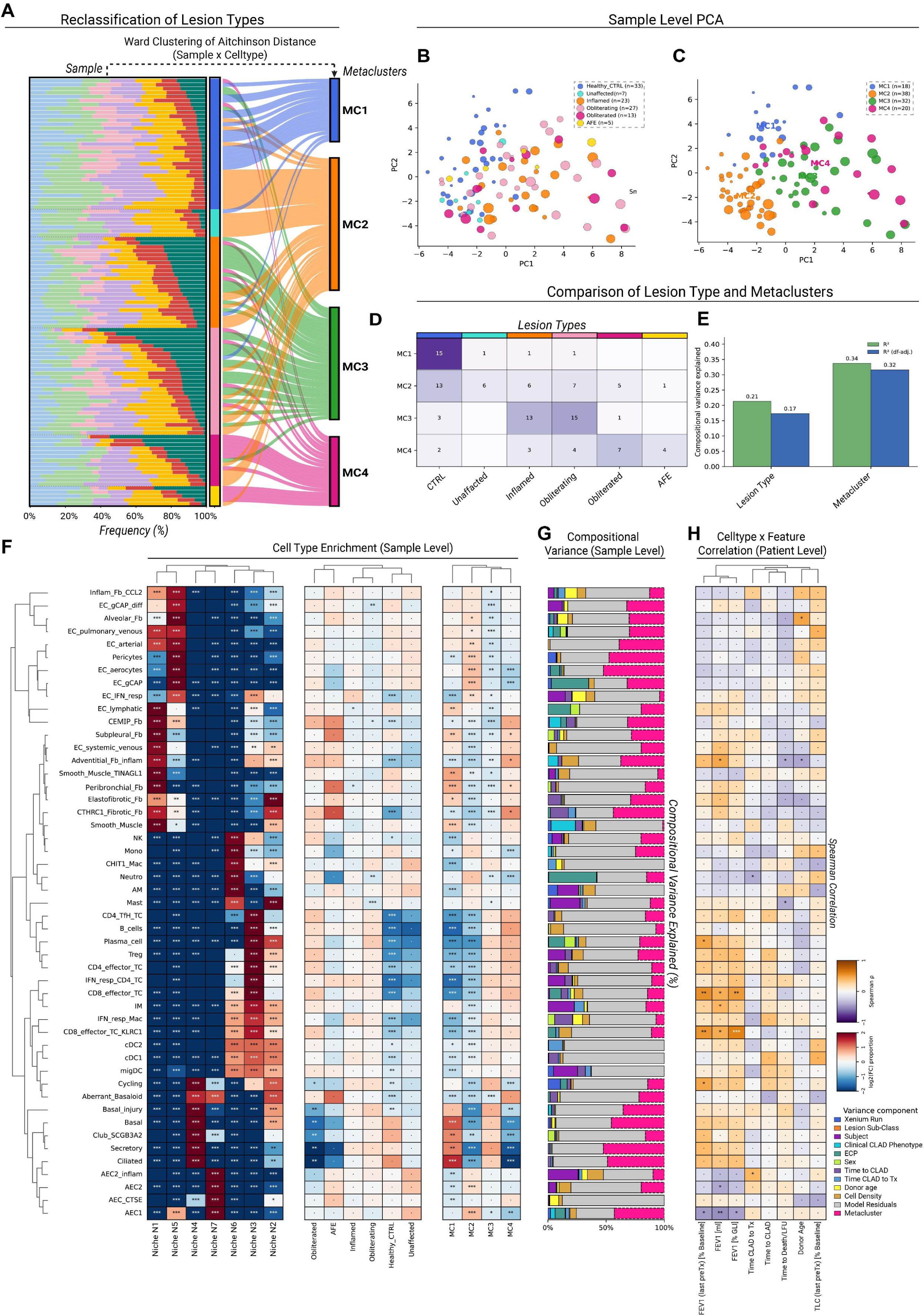
Unsupervised meta-clustering BOS samples. A) Stacked bar plots of relative niche frequencies per sample colored by niche and split by lesion sub-class. Subjects were ordered after frequencies of *Peribronchial Immune Niche* (N3). Sankey plots show transitions between histological lesion classification and unsupervised Aitchison-distance based metaclustering. B) Sample-level principal components (PC) colored by histological lesion sub-class. C) Sample-level principal components (PC) colored by metaclusters. D) Confusion matrix of histological lesion sub-classes and meta clusters. E) Bar plots of compositional variance explained (Aitchison R²) by lesion sub-classes versus metacluster, quantifying the fraction of total between-sample compositional variance attributable to each grouping. Values are shown raw (green) and degrees-of-freedom (df) corrected (blue) to account for the differing number of groups. F) Heatmaps show cell type enrichment across niches, histological lesion sub-classes and metaclusters with hierarchically clustered rows and columns. Enrichment is colored by log2 fold change in cell-type proportion. Significance was assessed by two-sided one-sample *Wilcoxon signed-rank tests* of per-sample log2 enrichment against the dataset-wide reference, with *Benjamini-Hochberg* correction applied independently within each panel. G) Stacked bar plots denote compositional variance explained per cell type. For each cell type, bars show the percentage of compositional variance attributable to each metadata and biological component, including the compositional metacluster (dashed) and unexplained residual variance. H) Exploratory spearman correlations were illustrated as heatmap with color coding of Spearman correlation coefficients (ρ) between per-sample cell-type proportions and clinical or demographic variables. P values remained unadjusted given the exploratory nature of this analysis.

## Discussion

In this study, we provide an atlas of BOS lesions combining snRNAseq with spatial transcriptomics, complemented by micro-CT-guided IF. Integration with our publicly available RAS data as a disease control allowed us to delineate both disease-specific and CLAD-overarching cellular principles of lung remodeling. We identified cellular hallmarks of fibrotic lung diseases in the BOS lung, including the "*Usual Fibrotic Niche*"^10^ composed of Aberrant Basaloid cells and *CTHRC1*+ fibrotic fibroblasts. Beyond these shared principles, we uncovered BOS-specific remodeling dynamics: an injury-associated basal cell state prone to differentiate into Aberrant Basaloid cells, depletion of peribronchial fibroblasts with local expansion of *CTHRC1*+ fibrotic fibroblasts, and a compositional shift in the peribronchial vasculature toward systemic venous EC at the expense of lymphatic EC. Surprisingly, this cell circuit was present not only “*Inside*” the remodeled airways, but also “*Outside*”, at the interface of broncho-vascular connective tissue and the alveolar parenchyma.

Deconstructing the BOS lesion into seven spatially defined tissue niches, we resolved two distinct tissue microenvironments at the luminal and alveolar interfaces of ongoing remodeling **(Fig.8A).** Notably, the epithelial–stromal circuit was not confined to the airway wall but extended "*Outside*," to the interface of broncho-vascular connective tissue and the alveolar parenchyma, revealing parenchymal involvement in a disease long considered primarily airway-centered. Within these niches, basal cells acquire an inflammatory, injury-associated state, and spatial transcriptomics allowed us to resolve them as a major reservoir of Aberrant Basaloid formation in human disease. Together with the immune infiltrates localizing to these niches, our data position the structural niches of the BOS lung as central to airway obliteration, rather than as a passive scaffold for immune-mediated injury.

**Figure 8.**
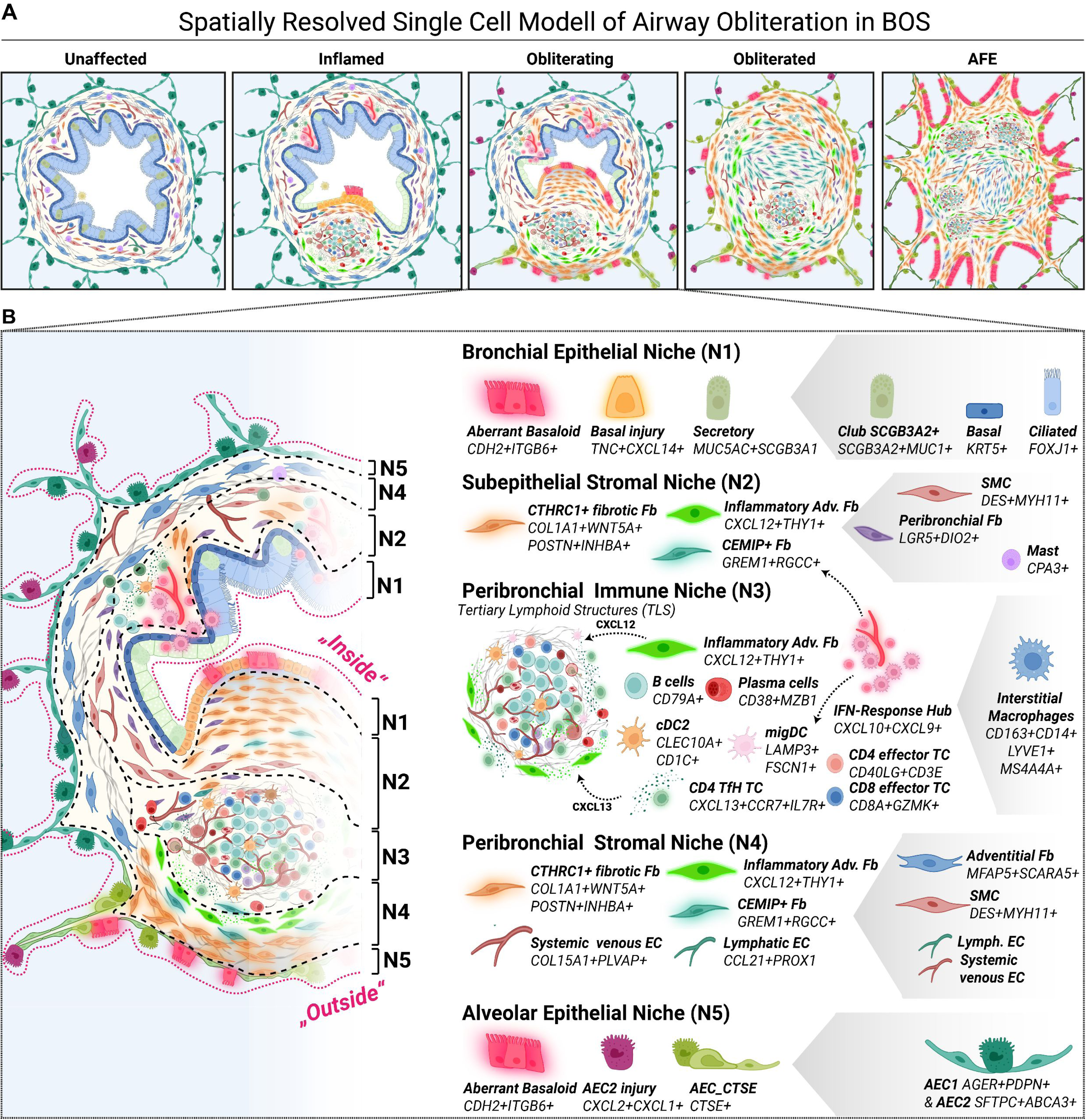
Summary of single cell-resolved stages in the obliterating BOS airway. A) Model of the distinct histological stages of airway obliteration in BOS. Alongside *Unaffected* airways, BOS lungs show the emergence of injury-associated basal cells and Aberrant Basaloid cells within the luminal epithelium already at the *Inflamed* stage, accompanied by the formation of early tertiary lymphoid structures (TLS) in the adjacent connective tissue. Airway obliteration (*Obliterating*) is mainly driven primarily by *CTHRC1*+ fibrotic fibroblasts, while fully *Obliterated* airways retain high numbers of these cells alongside an increasing accumulation of *CEMIP*+ fibroblasts. A subset of BOS lesions displays features of intra-alveolar fibroelastosis (*AFE*), with expansion of the lesional mesenchyme into the adjacent alveolar septa and dedifferentiation of the overlying alveolar epithelium into CTSE+ AECs and Aberrant Basaloid cells. B) Zoom-in into the cellular landscape of *Obliterating* BOS lesions. BOS niches of interest are delineated by dashed lines (N1-N5), and the two major sites of ongoing remodeling, "*Inside*" and "*Outside*", are marked by pink dashed lines. "*Inside*", bronchial epithelial cells acquire an injury-associated phenotype expressing *CXCL14* and *TNC* (*Bronchial Epithelial Niche*, N1), overlying *CTHRC1*+ fibrotic fibroblasts that arise in the *Subepithelial Stromal Niche* (N2) at the expense of resident peribronchial fibroblasts. In the *Peribronchial Immune Niche* (N3), TLS form, potentially in response to local *CXCL12* and *CXCL13* gradients secreted by inflammatory adventitial fibroblasts and CD4+ TfH-like cells, alongside scattered IFN-response hubs whose IFN-response macrophages infiltrate the N2 and N3 niches. Mirroring the stromal remodeling of N2, *CTHRC1*+ fibrotic fibroblasts and *CEMIP*+ fibroblasts also emerge in the outer *Peribronchial Stromal Niche* (N4), closely adjacent to the overlying *Alveolar Epithelial Niche* (N5) and coinciding with *CTSE*+ activation of the local AECs and their further dedifferentiation into Aberrant Basaloid cells, together recapitulating the "Usual Fibrotic Niche" observed across a range of fibrotic lung diseases. Created in BioRender. Schupp, J. (2027) https://BioRender.com/77x10o4

Building on previous phenotyping of the altered immune response in BOS^6–9^, our data indicate that the pathogenic cascade in BOS converges on a structural cellular landscape resembling the fibrotic parenchymal cell circuits of RAS^12^ and other ILDs^10,15,28–30^. This convergence extends to the vascular compartment: systemic venous EC have been shown to emerge in AFE in the context of RAS^12^ and PPFE^10^ and in other fibrotic lung diseases^15,31^. Collectively, our findings rationalize structural niche-directed treatment approaches and highlight structural cells as novel therapeutic targets in BOS.

CLAD, as an umbrella term, has traditionally spanned two main phenotypes: the obstructive BOS and the restrictive RAS phenotype^12^. The most recent ISHLT statement additionally defined an intermediate "*mixed*" phenotype, characterized by the co-presence of HRCT opacities and an obstructive PFT pattern^1^, which carries a poor survival comparable to RAS^32^. From a cellular perspective, our work challenges the current diagnostic paradigm of CLAD as three discrete phenotypes, as we observed the emergence of the same structural hallmark cells previously identified in RAS^12^ within the BOS lung as well. For BOS, it is established that patients with sequential involvement of both compartments exist, as BOS has been described to progress to a metachronous *mixed* phenotype. For RAS, in turn, the histopathological definition of its counterpart AFE includes prominent airway obliteration^11^, further underscoring the mutual similarities between BOS and RAS. Thus, we propose that CLAD represents a disease spectrum with shared, phenotype-overarching cellular drivers rather than an umbrella term for distinct post-transplant diseases.

Our study has limitations. First, as we analyzed end-stage CLAD specimens from re-transplantation, early drivers of disease initiation and progression may have been missed. Second, CTRL tissue originated from non-transplanted subjects, in part from peripheral tumor resections, and these subjects were significantly older than BOS patients, which may have introduced compositional confounding. Third, the spatial analysis relied on a pre-designed gene panel and focused on s=108 pre-selected regions of interest at distinct remodeling stages, favoring lesion-specificity over broader tissue context. Notably, the panel was informed by our unbiased snRNAseq data, with Xenium serving primarily for validation and localization.

## Conclusion

In summary, our study provides an unprecedented spatially resolved survey of the structural airway compartment throughout the histopathological sequence of airway obliteration in BOS, defining two major airway-centered sites of injury. The cellular similarities with RAS point to a shared disease spectrum and establish a molecular rationale for structural remodeling-directed treatment approaches in BOS and across the broader spectrum of CLAD. Finally, our BOS atlas offers an interactive, openly accessible resource to explore spatially resolved cell-specific gene expression changes in BOS, fostering mechanistic and translational research.

## Supplementary Material

### Supplementary Results

#### Supplementary description of the epithelial cell plasticity in BOS and RAS

Within the 62,935 curated epithelial nuclei we identified AEC1 (AEC1, *AGER, CAV1*), AEC2 (*SFTPC*, *FMO5*), basal cells (*KRT5*, *KRT17*), secretory cells (*MUC5B*, *MUC5AC*), and ciliated cells (*CFAP157*, *CFAP100,* **Fig. 2A, Table S2**). In addition, we uncovered four injury related epithelial cell states in the snRNAseq dataset namely Aberrant Basaloid cells (*MMP7*, *CDH2*, *KRT17*, *EPHB2*)^15^ *CTSE+* inflamed AEC1, and AEC2, that had been described across various interstitial lung diseases (ILD)^10,12,17^ as well as *CTSE*^low^ early inflamed AEC2 (*CXCL2*, *CSF3*, *CXCL1*, **Table S2**). Of these four cell states, we identified AEC1_CTSE (Padj._KW_=0.074) and Aberrant Basaloid cells (Padj._KW_=0.001) as two major epithelial hallmarks not only of the RAS but also of the BOS lung (**Fig. 2B, Table S9-10**). As described before^15^, Aberrant Basaloid cells expressed the full marker portfolio of genes related to terms of stemness (*EPHB2*, *MDK*), epithelial-to-mesenchymal transition (EMT; *CDH2*, *TNC*, *COL1A1*), basaloidness (*KRT17*, *TP63*), TGF-β activation (*TGFB2*, *ITGB6*, *ITGAV*), senescence (*CDKN1A*, *CDKN2A*), and others (*MMP7*, *GDF15*) across BOS and RAS (**Fig. 2C, Table S2**). This finding establishes Aberrant Basaloid cells as a well-conserved transcriptional cell state that is shared across fibrotic remodeling patterns, occurring not only in parenchyma-centered remodeling of RAS but also in airway-centered disease such as BOS.

#### Supplementary description of the intra-epithelial niches

*CellCharter* intrinsic stability analysis identified k=2 and k=4 as the most stable cluster numbers. Given the exploratory nature of this study, we profiled the cellular composition of the k=4 epithelial niches (E0–E3, **Fig.2E**). Niche E0 localized to the outer rim of the bronchovascular bundle, lining the peribronchovascular connective tissue, and showed a relative increase in Aberrant Basaloid and injury-associated basal cells versus CTRL (**Fig.2F**). Niche E1 represented the luminal bronchial epithelium, with the highest proportions of ciliated and secretory cells, and showed increased injury-associated basal cells, cycling epithelial cells and, to a lesser extent, secretory cells in BOS versus CTRL. The AEC1-(E2) and AEC2-centered (E3) niches reflected comparatively minor compositional shifts.

#### Supplementary description of the inflammatory fibroblast plasticity and gene programs

Hierarchical clustering resolved three inflammatory gene signatures (A–C) in inflammatory fibroblasts. Conserved signature A (*TNFRSF1A*, *TNFRSF12A*, *STAT3*) was expressed across *CCL2*+, *CCL2*- and subpleural fibroblasts (*WT1*+ *HAS1*+) and, to a lesser extent, in *CTHRC1*+ fibrotic fibroblasts, with expression increasing progressively from CTRL to BOS to RAS (**Fig.3C**, inflammatory signature A). Signature B (*CXCL2*, *SOD2*, *IL6*) was not regulated in CCL2-inflammatory adventitial fibroblasts but confined to their *CCL2*+ counterpart and to subpleural fibroblasts, the latter showing a comparable pro-inflammatory feature set in PPFE^10^. *CCL2*+ inflammatory fibroblasts remained transcriptionally stable across disease states (**Fig.3C**, inflammatory signature B), placing this subtype at the physiological end of the fibroblast plasticity spectrum. Signature C combined inflammatory (*CXCL12*, *DEPTOR*, *IL11*) and fibrotic (*COL1A1*, *IGF1*, *POSTN*) genes, induced in *CCL2*-inflammatory adventitial fibroblasts with an increasing trend across the disease continuum (**Fig.3C**, inflammatory signature C).

#### Supplementary description of the Mesenchymal Niches

*CellCharter*-based mesenchymal neighborhood clustering resolved four mesenchymal tissue niches (M0–M3, **Fig.3J–K**). Niche M0 represents the alveolar stromal niche, comprising predominantly pericytes and alveolar fibroblasts alongside a smaller population of *CCL2*+ inflammatory fibroblasts, thereby assigning this inflammatory fibroblast state to the alveolar compartment. Within M0, *CCL2*+ fibroblasts were relatively more abundant in BOS than in CTRL lungs (**Fig.3J**). Niche M1 encompasses the stromal niche of the loose connective tissue of the bronchovascular bundle surrounding the airways, dominated by adventitial fibroblasts (*MFAP5*, *PI16*, *SCARA5*). BOS-related remodeling of M1 was accompanied by increasing abundance of *CCL2*-adventitial inflammatory fibroblasts and *CEMIP*+ fibroblasts. The perivascular smooth muscle niche (M2), composed of *TINAGL*+ and *TINAGL*-smooth muscle cells (*DES*, *MYH11*), appeared largely unaltered in BOS. The fourth niche (M3) localized directly beneath the luminal airway epithelium, in close proximity to the aforementioned *CXCL14*+ injury-associated basal and Aberrant Basaloid cells. In CTRL airways, it comprised primarily peribronchial fibroblasts (**Fig.3J**), whereas BOS lesions showed a marked expansion of *CTHRC1*+ fibrotic fibroblasts, *CCL2*-inflammatory adventitial fibroblasts and, to a lesser extent, *CEMIP*+ fibroblasts, mirrored by a relative BOS-related depletion of peribronchial fibroblasts (**Fig.3J**).

#### Supplementary description of the Pan-Lineage Niches

*CellCharter* decomposition followed comprehensive lymphoid (**Fig.S10**) and myeloid (**Fig.S9**, **Table S11**) cell annotation, with k=7 clusters showing the highest compositional stability (**Fig.S6**).

##### *Bronchial Epithelial Niche* (N1)

EdgeR-based pseudobulk comparison between BOS and CTRL lesions, excluding *Unaffected* BOS lesions, confirmed upregulation of the basal injury markers TNC (blue) and CXCL14 (orange) and the emergence of Aberrant Basaloid-associated genes FN1, CDH2 and ITGB6 (**Fig.5C-D**). Among secretory cells, we noted increased expression of the Uroplakin-1b gene UPK1B (red, **Fig.5C**), previously reported to be induced in goblet cells in endobronchial biopsies of obstructive asthma^33^.

##### *Subepithelial Stromal Niche* (N2)

N2 spatially intersects N1 and the Peribronchial Stromal Niche (N4) and physiologically serves as the reservoir for peribronchial fibroblasts and, to a lesser extent, subepithelial smooth muscle cells (**Fig.5B**). In BOS, it became dominated by *CTHRC1*+ fibrotic fibroblasts forming focal protrusions, reflected by increased subepithelial *CTHRC1* (red), *GREM1* (yellow) and *MMP11* (orange) transcripts, with these protrusions populated by injury-associated basal and Aberrant Basaloid cells (**Fig.5C**).

##### *Peribronchial Immune Niche* (N3)

The immune aggregates of N3 resembled early-stage (TLS)^23–25^ and comprised a diverse set of TLS-associated immune cells such as B-cells (BC; *CD19*, *CD79A*, *MS4A1*), *CXCL13*-expressing T cells with a CD4+ T follicular helper-like phenotype (TfH-TC; *CCR7*, *IL7R*), CD8+ effector T cells (*CD8A*, *CD8B*, *GZMK*), cDC2 (*CD1C*, *CLEC10A*, *FCER1A*), and *LAMP3*+ mature migratory dendritic cells (migDC, *LAMP3*, *CCR7*, *FSCN1*). N3 further contained *SELP*+ systemic venous EC, which may have served as the local primary recruitment site. Relative frequency and density analyses showed increases in CD4+ TfH-like TCs, BCs, migDCs and cDC2, particularly in *Inflamed* and *Obliterating* BOS lesions (**Fig.S8**, **Table S25-28**). Spatial proximity between N3 and N4 was lower within BOS lesions than in CTRL airways (**Fig.5E**).

##### *Peribronchial Stromal Niche* (N4)

N4 usually engulfed the early forming TLS of N3 and showed high concentrations of inflammatory-fibroblast-derived *CXCL12* (dark green, **Fig.5C**), known to promote TLS formation^10,26,27^. Inflammatory adventitial fibroblasts were more abundant in remodeling-associated BOS lesions (**Fig.5B**), accompanied by relative shifts towards *CTHRC1*+ fibrotic fibroblasts and upregulation of the profibrotic transcription factors *RUNX1* and *RUNX2* and a set of ECM-remodeling enzymes (**Fig.5D**), consistent with a fibrotic transformation resembling the stromal alterations of the subepithelial airway compartment (**Fig.5C**).

##### *Alveolar niches* (N5–N7)

In BOS, the Alveolar Epithelial Niche (N5) was dominated by *CTSE*+ and *ITGB6*+ AECs with strongly increased Aberrant Basaloid cells (**Fig.5B**-**C**). The Alveolar Immune (N6) and Alveolar Interstitial (N7) niches were enriched for *CHIT1*+ interstitial macrophages (*CHIT1*, *TREM2*, *CD68*), alveolar macrophages (*MARCO*, *CD68*) and *CCL2*+ inflammatory fibroblasts relative to *CTRL* (**Fig.5B**-**C**).

#### Supplementary description on spatial LR analysis

Spatially informed ligand-receptor (LR) analysis (LIANA+, 389-gene Xenium panel, **Fig. S12A**) revealed elevated spatial co-expression of ligand-receptor-pairs (Moran’s R LR interactions) indicating increased cell signaling in BOS versus CTRL (**Fig. S12B**), increasing across *Inflamed*, *Obliterating*, *Obliterated*, and *AFE* lesions (**Fig. S12C**). Ward clustering of LR pseudobulk expression resolved three programs, one of which was induced across BOS lesions (**Fig. S12D**) and comprised three key interactions (**Fig. S12E**). First, focal allo-immune signaling was mediated by *CD28* and *CTLA4* on lesional T cells (CD4 TfH-like TCs, CD4 effector T cells and regulatory T cells (Tregs)) and by *CD80* and *CD86* on antigen-presenting cells (migDCs, cDC2, cDC1) (**Fig.S12F–I**), molecules previously implicated in chronic lung allograft rejection^34,35^. Second, induced *CXCL9*/*10*–*ACKR1* co-expression mapped back to pulmonary venous EC and IFN-response macrophages (**Fig.S12J**), consistent with augmented transendothelial chemokine presentation to circulating immune cells^36,37^ that may facilitate immune cell recruitment. Third, the *CD79A*–*FN1* interaction linked *CD79*+ immune cells such as plasma cells and B cells (**Fig.S12K**) to ECM-secreting fibroblasts such as *CTHRC1*+ fibrotic fibroblasts and inflammatory adventitial fibroblasts, an immune-mesenchymal crosstalk also observed by our group in PPFE-associated AFE^10^. Notably, systemic and pulmonary venous EC co-occurred within the same vessel (**Fig.4J**), suggesting a potential pulmonary-to-systemic phenotype conversion.

#### Supplementary description on meta clusters

*CTRL* airways mapped mostly to MC1, all histological lesion sub-classes to MC2, most *Inflamed* and *Obliterating* lesions to MC3, and most *AFE* and *Obliterated* lesions to MC4 (**Fig.7D**). MC1 represented the homeostatic airway (lowest immune cells, highest canonical basal, secretory and ciliated cells), MC2 alveolar-parenchyma-rich samples (highest AECs, aerocytes, pericytes, alveolar fibroblasts, and CCL2+ inflammatory fibroblasts), MC3 airway inflammation (TLS-associated cell types and injury-associated basal cells), and MC4 advanced fibrotic obliteration (highest *CTHRC1*+ fibrotic and inflammatory adventitial fibroblasts, Aberrant Basaloid cells, BCs, CD4+ TfH-like Tcells and plasma cells, with lower IFN-response cells) (**Fig.7F**).

## Supplementary Materials and Methods

### Sample selection

Bio-banked lung explant samples from 33 BOS patients were used for snRNAseq (s=14) and spatial transcriptomics (s=75, **Fig.1, Table 1**). These were complemented by s=46 CTRL specimens from n=33 subjects (**Table 1**), obtained either by peripheral tumor resection (n=8) or from downsized donor lungs (n=25), covering snRNAseq (s=13) and spatial transcriptomics (s=33, **Fig.S1**). Of these, snRNAseq data from s=13 samples derived from n=13 CTRLs had been published previously by our group (GSE284081^10^, s=4; GSE301982, s=9^12^). All tissue specimens were reviewed by a board-certified pathologist with more than 15 years of experience in lung transplantation (LTx). In addition, snRNAseq data from n=15 RAS patients (GSE301982)^12^ served as a disease comparator within the snRNAseq analysis. Selected formalin-fixed paraffin-embedded (FFPE) blocks were cut on routine microtome into 50µm thick sections for snRNAseq (three sections per sample), and to 5µm thick sections for spatial transcriptomics as well as to 8µm sections for IF analysis. Additional serially sectioned 5µm sections were used for Hematoxylin-Eosin and *Elastica van Gieson* staining. Slides were stored at room temperature for further use.

### Single Nucleus RNA sequencing (snRNAseq)

#### Nuclei isolation from FFPE specimens for 10X chromium fixed RNA profiling for multiplexed samples

Three 50 µm sections per FFPE block were transferred into Miltenyi C-tubes (#130-096-334). Paraffin was removed with three 10-min xylene washes (≥1 mL each), followed by rehydration in a 1-min graded ethanol series (100%, 70%, 50%, 30%) and three washes in 1X PBS with 0.5 mM CaCl2. After removal of excess liquid, 2 mL pre-warmed (37°C) Tissue Digestion Buffer (TDB) were added and samples digested on a gentleMACS Octo Dissociator (37°C, 45 min, with 30-s clockwise and counterclockwise spins at 2,000 rpm). C-tubes were centrifuged (300 rcf, 30 s) and pellets resuspended in their supernatant. Suspensions were passed through 70 µm strainers (PluriSelect #43-50070-51), rinsed with PBS, centrifuged (850 rcf, 5 min, 4°C), resuspended in 1 mL chilled Quenching Buffer and filtered through 20 µm strainers (PluriSelect #43-10020-50). Nuclei concentration and viability were measured on a LUNA-FL Dual Fluorescence Cell Counter using AO/PI staining. For fixation, 0.1 volume of pre-warmed Enhancer (Chromium Next GEM RNA Profiling Sample Fixation Kit) was added, and samples were stored at 4°C for up to 1 week before Fluorescence activated nuclei sorting (FANS).

#### Fluorescence activated nuclei sorting (FANS) of DAPI+ single nuclei

Nuclei were sorted at the MHH Cell Sorting Research Facility. Immediately before sorting, nuclei were stained with DAPI (Thermo Scientific, final 1–10 µg/mL, adjusted per sample). Sorting used a 100 µm nozzle at 35 psi (FACSAria III Fusion, FACSAria IIu, BD, or MoFlo XDP, Beckman-Coulter). Intact single nuclei were gated by light-scatter properties and DAPI-based DNA content, with a narrow DAPI gate excluding fractional-DNA nuclei and debris and scatter parameters removing doublets. 5×10⁵ DAPI+ single nuclei per specimen were sorted into 15-mL tubes containing 1 mL 0.5X PBS + 0.02% BSA. Reanalysis of 300 sorted nuclei per specimen confirmed a homogeneous population. Sorted nuclei were kept on ice and processed in batches of 16 specimens.

#### 10x genomics chromium fixed RNA profiling for multiplexed samples

Sorted DAPI+ nuclei were processed per the *10x User Guide CG000527 Rev B* with the following choices. Experiments were designed to maximize number of cells without sub-pooling, using a post-hybridization pooled-wash workflow, with hybridization in PCR strips. Because 500,000 nuclei were sorted per specimen, the first post-hybridization counting step was omitted and all 16 specimens per pool were fully pooled, with near-identical nuclei numbers ensured by FANS. The pooled pellet was resuspended in 25% of the protocol volume to reach a high concentration before 20 µm filtration (pluriStrainer Mini 20 µm). For "Targeted Cell Recovery" (128,000 nuclei), a 20% larger stock volume than calculated was used and brought to 40 µL with Post-Hyb Resuspension Buffer. Sample Index PCR used 12 cycles.

#### RNA library quality control

After construction, each library underwent three QC steps. A 1:20 dilution in Buffer EB (Qiagen) was quantified by Qubit (Qubit 1X dsDNA HS/BR Kit, Qubit 4 Fluorometer, Invitrogen). 5 µL of each diluted library was sent to the MHH Research Core Unit Genomics (RCUG) for fragment-length analysis (HS) and Qubit quantification. Each library was additionally quantified by qPCR (KAPA Library Quantification Kit for Illumina, Roche KR0405 v11.20). Stock and diluted libraries were stored at -20°C.

#### Sequencing

Libraries were sequenced on an Illumina NovaSeq6000 (Institute of Human Genetics, MHH) with 28 bp Read1, 90 bp Read2 and 10 bp for i5/i7. Base calls were demultiplexed to FASTQ with Cell Ranger (v7.1.0) *mkfastq*.

#### Data processing

Reads were processed with Cell Ranger (v7.1.0) and aligned to GRCh38 (GENCODE v32/Ensembl 98, GRCh38-2020-A) using the Chromium Human Transcriptome Probe Set v1.0.1. Count matrices were analyzed with Seurat (v4.3.0.1) in R (v4.2.1). UMI counts were normalized to 10,000 UMIs per cell and natural-log transformed with a pseudocount of 1^15,16,38^. Quality control metrics are provided in **Table S29**.

#### Low-quality filtering

Mitochondrial content was calculated per nucleus from "MT-" genes. Quality filtering was performed iteratively per sample after normalization, scaling, dimensionality reduction and lineage-marker plotting to minimize information loss. Nuclei with <100 counts, <100 features or >15% mitochondrial reads were removed. Pre-cleaned samples were then iteratively integrated and clustered by reciprocal PCA (RPCA), using *FindVariableGenes*() (nFeatures=2000), *FindIntegrationAnchors*(), *IntegrateData*() and *ScaleData*()^16^.

#### Integration, dimensionality reduction und clustering

Variance-selected principal components were used to compute Euclidean distances and a nearest-neighbor graph, followed by Louvain clustering and UMAP visualization. Clusters without significant differential expression were merged iteratively until all nuclei across subjects were assigned to consistent cell types. Lineages (Epithelium, Mesenchyme, Endothelium, Lymphoid, Myeloid) were defined by marker expression (Fig. S3A). Each lineage was re-embedded and re-clustered at least twice to remove multiplets, empty cells and debris, then re-integrated by RPCA with final dimensionality reduction and graph embedding as above.

#### Clustering

*FindNeighbours*() was run with default k=20 and Euclidean distance. Cluster resolution was increased until no major inter-cluster differential expression remained.

#### Cell type annotation

Cell type markers were identified with Seurat’s *FindMarkers*() (*Wilcoxon rank-sum*, each cluster vs. all others, *Bonferroni*-adjusted P<0.05). Markers were ranked by diagnostic odds ratio (DOR, positive expression X>0, pseudocount 0.5). Cell types were annotated in a supervised manner (by logDOR) together with established literature markers ^14–16,19,39^.

#### Pseudotime analysis

Trajectory inference was performed with *Palantir* (v1.4.4) in *Python* (v3.13.9), invoked from R (4.2.1) through the SCP wrapper *RunPalantir*, which converts the Seurat object to an *AnnData* object via reticulate. The RNA assay was used as expression input, with principal component analysis as the linear reduction and the 30-PC UMAP embedding as the nonlinear reduction. Cells were grouped by fine-grained epithelial cell type annotation. Aberrant Basaloid cells were specified as the early group, and the representative cell of this group was used directly as the trajectory start without further repositioning. AEC1 and AEC2 were defined as terminal groups, with terminal cells likewise taken as assigned. Pseudotime and branch probabilities were computed over 1,200 waypoints with a maximum of 25 iterations. Finally, the returned pseudotime was inverted (1 − pseudotime), so that increasing values denote progression toward the Aberrant Basaloid endpoint. Pseudotime and differentiation potential (entropy) were transferred back to the Seurat object by matching cell barcodes.Downstream analyses were restricted to the terminal segment of the trajectory (inverted pseudotime > 0.7), capturing the transition zone between injured alveolar epithelial states and Aberrant Basaloid cells. Genes with pseudotime-dependent expression were identified with *RunDynamicFeatures* using 200 candidate features and a fixed random seed. Expression dynamics of selected hallmark genes were visualised as smoothed fits with confidence bands along the pseudotime axis after z-score scaling, stratified by cell type and by disease group.

#### Subject-level ordinal gene trend analysis across diseases

Trend testing was performed on the subject-collapsed log-normalized pseudobulk matrix. Analyses were run independently per cell type. For each gene, the association with the ordered disease axis was quantified by Spearman rank correlation between per-subject expression and the ordinal numeric disease code, with significance assessed by an asymptotic correlation test. Group means and pairwise deltas (CTRL vs. BOS, BOS vs. RAS, CTRL vs. RAS) were computed alongside, and a gene was defined as monotonically increasing when mean expression rose across both consecutive transitions. P values were adjusted within each cell type using the *Benjamini-Hochberg* method. Genes were called as showing a significant disease-associated increase when they combined an adjusted P<0.05, a positive correlation coefficient, and monotonicity of group means. Results were visualized as per-cell type scatter plots of Spearman rho against −log10(Padj.). Top-ranked genes of interest were annotated manually into three functional programs (“*Migration*”, “*Inflammation*”, “*Fibrosis*”) for labelling.

#### Pseudobulk differential gene expression across disease states

Differential expression between disease states was assessed per cell type using a pseudobulk approach as follows: raw counts of the RNA assay were summed across all nuclei of a given cell type within each subject, yielding one expression profile per cell type and subject. Pseudobulk profiles derived from fewer than 10 nuclei were discarded, and a cell type was tested only if at least three subjects per disease group remained after filtering. Lowly expressed genes were removed with *edgeR’s filterByExpr()* function, and libraries were normalized by the trimmed mean of M-values method. Dispersions were estimated with robust empirical Bayes shrinkage, and a quasi-likelihood negative binomial generalized linear model was fitted with disease state as the only covariate (*design ∼0 + disease*). Contrasts were defined as BOS versus CTRL and BOS versus RAS and evaluated by quasi-likelihood F-test. P values were adjusted for multiple testing across all tested genes of a given cell type and contrast using the *Benjamini-Hochberg* procedure. Genes were considered differentially expressed at an adjusted p-value below 0.05, an absolute log2 fold change above 0.5 and an average abundance above 1 log2 counts per million. For each gene, the fraction of expressing nuclei per group and the log diagnostic odds ratio with Haldane-Anscombe correction were reported as complementary effect size measures. Analyses were performed with *edgeR* (v.4.3.1) in R (v.4.2.1).

### 10X Genomics Xenium Spatial Transcriptomics

#### Assembly of tissue multiarrays (TMAs)

For tissue microarray generation, BOS lesions and CTRL airways annotated on hematoxylin and eosin-stained sections were punched from the corresponding FFPE blocks using a custom 3×3 mm square punch biopsy device. Punched fragments were re-embedded into pre-warmed paraffin in a 3×6 (3×7) array format, yielding 5×18 specimens and 1×21 specimen per block (**Fig.S1**). To preserve the orientation of the fragments for downstream processing, a sacrificial tissue piece was included in one corner of the new array.

#### 10X Genomics Xenium sample preparation

Spatial transcriptomics was performed on the Xenium Analyzer (10x Genomics) using v1 chemistry together with the Cell Segmentation Staining workflow, following the manufacturer’s user guides for deparaffinization and decrosslinking (CG000580), in situ gene expression including cell segmentation (CG000479, CG000749) and instrument operation (CG000584) without modification. 5µm thick FFPE sections mounted on Xenium slides were baked for 2 h at 60 °C, equilibrated to room temperature for 7 min and deparaffinized in xylene (2 × 10 min) followed by graded ethanol (2 × 3 min 100%, 2 × 3 min 96%, 3 min 70%) and nuclease-free water (20 s). Slides were assembled into Xenium cassettes and decrosslinked in Xenium FFPE Tissue Enhancer supplemented with 8 M urea and diluted Perm Enzyme B for 30 min at 80 °C followed by 10 min at 22 °C, then washed three times in PBS-T. Pre-designed gene expression probes and add-on custom probes (289 + 100 custom targets in total) were denatured for 2 min at 95 °C, chilled on ice for 1 min and combined with Xenium Probe Hybridization Buffer and TE buffer. Hybridization proceeded for 17 h on the thermal cycler using the manufacturer’s default program.

On the following day, samples were washed in PBS-T and Xenium Post Hybridization Wash Buffer, ligated for 2 h in Xenium Ligation Buffer containing Ligation Enzymes A and B, and amplified for 2 h in Xenium Amplification Mix supplemented with Amplification Enzyme, with PBS-T washes between steps and TE buffer after amplification. For cell segmentation staining, sections were dehydrated and rehydrated through four sequential 2-min ethanol washes (70%, 100%, 100%, 70%), washed in PBS-T and blocked in 1× diluted Xenium Block and Stain Buffer for 1 h at room temperature. Xenium Multi-Tissue Stain Mix was reconstituted in 220 µl of the same buffer, incubated for 30 min at room temperature and cleared by centrifugation for 10 min at 14,000 rcf and 4 °C. Then 100 µl of the supernatant was dispensed into the Xenium Cassette Insert, the well was sealed and staining proceeded for 21 h at 4 °C.

Inserts were floated off in PBS-T and removed with forceps, and sections were incubated in resuspended Xenium Stain Enhancer for 20 min at room temperature. Autofluorescence was quenched by 10 min in diluted Reducing Agent B, sequential 70% and 100% ethanol washes and 10 min in Autofluorescence Solution in the dark, followed by three 2-min washes in 100% ethanol, drying for 5 min at 37 °C and rehydration in 1× PBS and PBS-T. Nuclei were stained for 1 min in Xenium Nuclei Staining Buffer in the dark and washed three times in PBS-T. Decoding Reagent Modules A and B and the Cell Segmentation Detection Module were equilibrated as specified, mixed by gentle inversion and centrifuged for 1 min at 300 rcf. Instrument Wash Buffer, Sample Wash Buffers A and B and Probe Removal Buffer were prepared fresh, degassed for 30 min at room temperature and loaded together with cassettes, reagent plates and consumables. Regions of interest were selected after the 1-h sample scan, and the run was executed with default instrument settings.

#### 10X Genomics Xenium gene panel selection

The final panel comprised 389 unique genes, of which 289 were derived from the Xenium Human Lung Panel v1 (1000601) and 100 from a custom add-on panel (<u>1000645, Panel ID: 8T49U3</u>, **Table S30**). Custom targets were selected from the snRNAseq analysis described above by ranking the top 100 differentially expressed genes per cell type and retaining those most informative for cell type discrimination in a supervised manner.

#### 10X Genomics Xenium cell segmentation

Cells were segmented using the onboard segmentation algorithm of the Xenium Analyzer (version 4.0, 10x Genomics). Nuclei were first delineated from the DAPI morphology image, and adjacent nuclear boundaries were resolved into non-overlapping objects. Cell outlines were then defined using the multi-tissue cell boundary stain. Cell identity was assigned on the basis of nuclear segmentation alone.

#### Virtual sample separation, quality filtering and data preprocessing

Tissue coordinates of each sample were extracted by drawing manual region of interest in the Xenium Explorer (v4), followed by export of the coordinates as csv. Xenium output bundles were converted to *SpatialData* objects and stored as *Zarr* archives. Regions of interest corresponding to individual tissue microarray cores were defined from the instrument selection coordinates and extracted by bounding-box query in the global coordinate system. Each cropped object was written out in its raw state. 2 non-BOS / non-CTRL samples and 1 detached CTRL sample were excluded from downstream analysis yielding a total of s=108 samples. Per-sample objects were concatenated into a single dataset. Quality control followed recently published best practices guidelines^40^ and cells with fewer than 10 transcripts were excluded. No gene-level filtering was applied given the targeted nature of the panel. Raw counts were retained as a separate layer, normalized to a target sum of 100 transcripts per cell, log1p-transformed, stored as a log-normalized layer and scaled without centering to a maximum value of 10. Principal component analysis was computed on 50 components, and batch effects between samples were corrected with Harmony. The corrected embedding served as input for cosine-distance nearest-neighbor graph construction and UMAP embedding, both computed with the GPU-accelerated *rapids-single-cell* implementation.

#### 10X Genomics Xenium tissue surface calculation

Tissue surface area per sample was approximated from the segmented cell centroids. For each specimen, centroid coordinates were extracted from the spatial embedding, and the convex hull of the resulting point set was computed. The enclosed polygon area, reported in µm^2^ served as the sample surface estimate and as the denominator for all cell density calculations (cells per mm²).

#### Dimensionality reduction, clustering and cell type annotation

Lineage-level objects were subclustered iteratively to resolve cell states. For each iteration, principal component analysis was recomputed on 50 components, samples were integrated with Harmony, and a nearest-neighbor graph was built on the corrected embedding using cosine distances, followed by UMAP embedding and Leiden clustering. Cluster-defining genes were identified by Wilcoxon rank-sum testing. Clusters expressing markers of other lineages or hallmarks of segmentation-derived “doublets” were removed, and the remaining cells were re-processed from raw counts under the same workflow. This cycle was repeated until all clusters were lineage consistent. Neighborhood and clustering parameters were adjusted across iterations to match the decreasing dataset size and increasing transcriptional similarity of the retained cells. After the final iteration marker genes per sample were calculated by Bonferroni adjusted Wilcoxon rank sum test analogously to computations for cell type markers within the snRNAseq pipeline. Clusters were then annotated manually based on canonical marker gene expression. All computations used the GPU-accelerated *rapids-singlecell* implementation of the *scanpy* workflow, with fixed random seeds.

#### Cross-platform validation of cell-type annotation

Cell type annotations derived from spatial transcriptomics were validated against the snRNAseq reference by comparing transcriptional profiles across platforms. For this purpose, the integrated Seurat object was exported to *h5ad* format via *SeuratDisk* and read into Python as an *AnnData* object, with the raw count matrix reattached from the .raw slot and verified against the log-normalized expression matrix by comparing their nonzero patterns. Given the scope of this study, the analysis was restricted to structural cell types and to genes measurable on both platforms. Discriminating genes were defined on the Xenium data by *rank_genes_groups* (*Wilcoxon rank-sum test*) per cell type, retaining genes with a log2 fold change ≥0.25 and an adjusted P ≤0.05. The 20 top-ranked qualifying genes per cell type were pooled into a common feature space. For each platform, mean log-normalized expression of these genes was computed per cell type and each gene was z-scored across cell types within its platform, so that the comparison reflects cell type specificity rather than platform-dependent capture efficiency. Transcriptional similarity between every Xenium and every Flex cell type was then quantified by Spearman rank correlation across the shared feature space and displayed as a heatmap with cell types grouped by lineage on both axes.

#### Assessment of cell densities and proportions

Cell densities and cell type proportions were analyzed per sample, with the sample treated as the unit of analysis. For densities, tissue area was approximated as described above and converted to mm², and densities were calculated as the number of cells of a given cell type or niche divided by area. For proportions, the relative frequency of each cell type was computed per sample as its fraction per investigated lineage. Both measures were compared across lesion sub-classes per cell type by *Kruskal-Wallis* test, with *Benjamini-Hochberg* adjustment across all cell types tested. Cell types not reaching significance in the omnibus test were excluded from *post-hoc* analysis. *Post-hoc* comparisons of all lesion sub-class pairs were performed by *Wilcoxon rank-sum test* with *Benjamini-Hochberg* adjustment within each cell type. Adjusted P-values<0.05 were considered statistically significant.

#### Niche identification

Spatial niches were defined with *CellCharter*^18^. For each sample, a spatial neighborhood graph was constructed on cell centroids by Delaunay triangulation. Neighborhood-aware representations were generated by aggregating the Harmony-corrected principal component embedding over three concentric layers of neighbors. Niche assignment used a Gaussian mixture model, and the number of clusters was selected by repeated fitting across k = 2 to 10. The most stable fitting (k=7) was used for pan-lineage niche annotation. For niche analysis within the structural niches we pursued a trade-off of stability and biological/anatomical interpretability. Cell type enrichment per niche was expressed as the log2 ratio observed to expected cell counts, the latter derived from the global cell type frequencies scaled to niche size. Disease-related shifts were quantified by computing, for each sample, the percentage each cell type contributed to a given niche, averaging these per-sample percentages within disease groups and expressing the BOS-versus-CTRL difference as a log2 ratio. Spatial relationships between niches were assessed by *CellCharter’s* inbuilt neighborhood enrichment, computed separately for BOS and CTRL samples. Differential neighborhood enrichment between conditions was tested by permutation with sample as the library key (100 permutations), and pairs with P<0.05 were considered significant.

#### Niche surface estimation

Niche expansion was quantified by tissue surface rather than cell number, so that it reflects the area a niche occupies independently of local cell density. Each section was partitioned into a grid of 50×50 µm bins, and every occupied bin was assigned the niche of the majority of its cells. Niche surface per sample was computed as the number of assigned bins multiplied by the bin area. Binning was preferred over a convex hull estimate because niches are interdigitated and frequently non-convex, so that a hull would overestimate their extent and assign overlapping areas to different niches. For the broncho-vascular compartment, airway-associated niches were renormalized within each sample to sum to 100%, expressing each as its share of the bundle. Relative surfaces were compared across lesion sub-classes by *Kruskal-Wallis* test, followed by pairwise *Wilcoxon-Rank sum tests*. P values were adjusted by the *Benjamini-Hochberg* procedure within each niche for the pairwise comparisons.

#### Longitudinal frequency analysis

Treating lesion sub-classes as equidistant units of increasing severity (*CTRL*, *Unaffected*, *Inflamed*, *Obliterating*, *Obliterated*, *AFE*), cell type frequencies were computed per sample within each niche as the proportion of a given cell type among all cells of that niche, so that each sample contributes one observation per niche and cell type. Trajectories were modelled by weighted least-squares polynomial regression against this severity axis, weighting each sample by its cell number in the niche. The polynomial degree was selected per curve as the lowest degree from linear to cubic without significant lack of fit (F-test, P≥0.05), falling back to the corrected Akaike information criterion where no degree qualified, and constrained to remain identifiable given the available samples and lesion sub-classes. Fitted mean curves are displayed with 95% confidence bands.

#### Cell type proximity analysis

Co-localization networks were inferred from spatial proximity to Aberrant Basaloid cells, B cells and IFN-response macrophages, excluding BOS samples annotated as *Unaffected*. For each partner cell, the mean Euclidean distance to its ten nearest target cells was computed by *BallTree* search within samples and summarized as the median per cell type and disease group. Networks were rendered as lineage-ordered circos plots in which node size encodes spatial density (cells per sample surface, scaled to the 99th percentile), edge width and opacity encode median nearest-neighbor distance on a gamma-corrected scale shared across conditions.

#### Niche-resolved differential expression

To preserve the sample as the unit of analysis, raw Xenium counts were summed across all cells of a given niche within each sample, yielding one pseudobulk profile per niche and sample. Niche-sample combinations with fewer than 30 cells were discarded. Pseudobulk matrices were transferred to R via *rpy2* and *anndata2ri* and analyzed with *edgeR*. After removal of lowly expressed genes and library size normalization, an intercept-free design was fitted over the combined niche*lesion-class factor, followed by dispersion estimation and quasi-likelihood negative binomial model fitting. Every lesion sub-class was contrasted against *CTRL* within the same niche, so that all comparisons remain internal to a niche, and contrasts were evaluated only when both sides comprised at least three pseudobulk samples. P values were corrected using the *Benjamini-Hochberg* procedure.

#### Spatial L–R analysis

Cell-cell communication was inferred with *LIANA+* on log-normalised Xenium expression using the consensus ligand-receptor resource restricted to the 389-gene panel. Cluster-level interactions between cell types were scored by the *rank_aggregate* consensus of *CellPhoneDB*, *Connectome*, *log2FC*, *NATMI* and *SingleCellSignalR*, requiring expression in at least 10% of cells per cluster and assessing specificity by permutation, with a specificity rank ≤0.05 considered specific. The analysis was performed on the full dataset and repeated within disease groups and within lesion sub-classes, excluding classes with fewer than 500 cells. Selected signaling axes were visualized as chord diagrams, in which rim sectors represent cell types and ribbons individual source-to-target interactions. Ribbon width scales with interaction strength (-log10 magnitude rank) and ribbon color denotes either the ligand-receptor pair or the sending cell type, with arrowheads marking the receptor end. Spatially informed scores were computed with *LIANA+* bivariate. Neighborhood graphs were built per sample using a Gaussian kernel (bandwidth 200 µm, cutoff 0.1), avoiding neighborhoods to cross sample border, with the bandwidth selected empirically from the neighbour-count profile of a representative section. Local co-expression was quantified by spatially weighted cosine similarity and summarized per section by bivariate Moran’s R. Per-section Moran’s R was averaged across significant interactions and compared between disease groups by Wilcoxon rank sum test and across lesion sub-classes by *Kruskal-Wallis* with *Benjamini-Hochberg*-corrected pairwise comparisons. For the heatmaps, ligand-receptor pairs were retained when scored in at least 50% of sections and, for the stage-resolved analysis, when spatially significant in at least half of a class’s sections. Qualifying pairs were ranked by the variance of their mean Moran’s R across classes, prioritizing interactions whose spatial co-localization changes along the remodeling continuum. Scores were z-scored per pair and clustered by Ward linkage. For spatial plots of individual ligand-receptor pairs, spatial plots were colored by their local interaction score, with the color scale shared across the displayed sections of a given pair.

#### Meta-cluster inference

Samples were re-classified in an unsupervised manner as: cell type counts per sample were centered log-ratio transformed with a pseudo count of 0.5, and samples contributing fewer than 200 cells were excluded. Samples were clustered by Ward linkage on *Euclidean* distances in centered log-ratio space, corresponding to the *Aitchison* distance, and the number of clusters was selected by maximizing the mean silhouette width across candidate solutions from k=2 to k=8. The resulting metaclusters were renumbered by their mean position along the histological remodeling continuum, so that MC1 corresponds to the least and the highest MC to the most advanced compositional state. To compare the histological and the data-driven classification, multivariate variance explained was computed in Aitchison space for both groupings as the between-group sum of squares divided by the total sum of squares, together with a degrees-of-freedom-penalized variant and a pseudo-F statistic. Significance of pseudo-F was assessed by 999 permutations of the group labels.

#### Cell type enrichment analysis

Cell type enrichment was quantified as the log2 ratio of the proportion of a given cell type within a group to its proportion across the whole dataset, with a pseudocount added to both terms to stabilize ratios for rare populations. Enrichment was computed in parallel for spatial niches, histological lesion sub-classes, and compositional metaclusters. Significance was assessed on per-sample proportions expressed against the same dataset-wide reference and tested against zero within each group by two-sided one-sample *Wilcoxon signed-rank* tests, retaining the sample as the unit of observation. Samples with fewer than 20 cells were excluded and tests required at least four samples per group. P values were corrected by the *Benjamini-Hochberg* procedure separately within each panel, so that niche, lesion-class, and metacluster comparisons form independent test families. Cell types and panel columns were ordered by hierarchical clustering (average linkage, *Euclidean* distance) on the enrichment values, using pairwise-complete distances.

#### Compositional Variance Analysis

Variance in centered log-ratio abundance was partitioned per cell type with *variancePartition*, called via *rpy2*. Categorical covariates (run, lesion sub-classes, subject, metacluster, clinical CLAD phenotype, azithromycin and extracorporeal photopheresis treatment, sex) entered as random effects, continuous covariates (time to CLAD, time from CLAD onset to re-transplantation, donor age, and cell density derived from the convex-hull area of the spatial coordinates) as fixed effects after z-scoring. Missing categorical values were retained as an explicit level, whereas samples with missing continuous values were excluded by complete-case filtering. Terms were dropped automatically when they exceeded 40% missing values, carried fewer than two levels, resolved to one level per sample, or were collinear with an earlier-listed continuous covariate (|r| > 0.7). Variance fractions, including the unexplained residual, are reported per cell type as percentages of total variance.

#### Exploratory correlations of per subject cell type frequencies and clinical variables

Associations between cell type composition and continuous clinical variables were computed per subject and correlated with one clinical value per patient by *Spearman* rank correlation. P values<0.05 were considered statistically significant. These correlations are reported with uncorrected P values and remain therefore exploratory.

### HRCT-Segmentation

Last HRCTs scans prior re-transplantation were analyzed using a deep learning-based software tool (AVIEW Lung Texture, Coreline Soft, Seoul, Korea). This system automatically segments the lung parenchyma and classifies it into six patterns: normal lung, reticulation, honeycombing, ground-glass opacity, consolidation, and emphysema.

### Micro-CT serial sectioned BOS lesions

Lungs of patients undergoing re-transplantation for end-stage BOS were air-inflated via cannulation of the main stem bronchus and fixed in liquid nitrogen fumes. The lungs were sliced from apex to base in 2 cm slices and cores were systematically removed and scanned with microCT (Flex CT voxel size of 15 µm). These microCT scans were used to identify typical BOS lesions. After fixation and FFPE embedding, these microCT scans were used to navigate to BOS lesions by capturing sections cut at 8 µm before and during airway obliteration.

### Immunofluorescence protein staining

FFPE section (2µm) were rehydrated through xylene and ethanol, and antigen retrieval was performed in 95°C water with 1% Antigen Unmasking Solution (Vector, H3301) for 20 min, followed by cooling to RT in 1X PBS. Nonspecific binding was blocked with 2.5% normal donkey or horse serum (Biozol JIM-017-000-121 or Vector S-2012, matched to the secondary antibody species) for 20 min. Incubation with the primary antibody (**Table S31**) was performed for 1 h at RT, with PBS washes between steps. For two-color panels, DyLight 488 anti-mouse and DyLight 594 anti-rabbit secondaries (VectaFluor Duet Kit, Vector DK-8828, **Table S31**) were applied for 1 h. For three-color panels, donkey anti-rabbit Rhodamine Red-X (Biozol JIM-711-295-152), anti-mouse AlexaFluor 647 (Life Technologies A32787) and anti-rat AlexaFluor 488 (A21208) were applied as a cocktail for 1h at RT, followed by a 1 h incubation with a directly conjugated primary antibody against an additional marker at RT. Autofluorescence was quenched with the Vector TrueView Kit (SP-8400-15), and slides were mounted in DAPI-containing antifade medium (Vector H-1800-10) and stored at 4°C in the dark.

### Immunofluorescence Microscopy

Whole slides were scanned on a Zeiss Axio Scan 7 with a 20x/0.8 Plan-Apochromat objective, using the 96 HE BFP 450/40, 38 eGFP 525/50, 43 HE DsRed 605/70, 26 AlexaFluor 660 685/50 and 50 Cy5 690/50 filter sets.

### Use of Generative AI

Generative artificial intelligence tools (ChatGPT, OpenAI; Claude, Anthropic) assisted with code development, debugging and refinement for data analysis and visualization, and with language refinement for clarity. All AI-assisted code and text were reviewed and validated by the authors, who performed and verified all analyses and scientific interpretations and take full responsibility for the manuscript.

### Statistical Analysis

Cell frequencies, spatial cell type densities, and niche surfaces were compared across lesion sub-classes and diseases using *Kruskal-Wallis* test. *Post hoc* inter-group comparisons were performed with the Wilcoxon rank-sum test, adjusted for multiple testing using the Benjamini-Hochberg method. Correlations between snRNAseq- and spatial transcriptomics-derived measures as well as across clinical metadata and cell type frequencies were assessed using Spearman’s rank correlation. P values <0.05 were considered statistically significant, with significance levels indicated as *: P≤0.05, **: P≤0.01, ***: P≤0.001 and ****: P≤0.0001. snRNAseq and clinical data analyses were computed in R (v4.2.1, The R Foundation), and spatial transcriptomics (10X Xenium) analyses in Python (v3.13.9).

### Web Tool

A web-based interactive application [https://breath.mh-hannover.de/BOSatlas.html] was developed using Shiny for Python to facilitate exploration and visualization of the data. The application provides an interactive interface for querying the processed datasets and generating visualizations of cell populations, gene expression, co-expression patterns, and other relevant analyses. It was deployed on a Linux server. The application and its deployment configuration were containerized to ensure reproducibility and portability.

## Supporting information

Supplementary_TablesS1_to_S31

Supplementary_FiguresS1_to_S12

## Acknowledgments

The authors thank the Research Core Unit for Laser Microscopy at Hannover Medical School for their support. The authors thank all study participants for their permission to use their respective tissue specimens for research.

## Authors contributions

JCS, JCK, CF, and JG conceived and supervised the project. JCS acquired funding. LC, LML, LG and MB performed nuclei isolation, barcoding and library generation. RE, JCK and LN assembled the TMAs. LC, AB, WR, JR, LG, RE, CP and EC generated the Xenium data. JR, HY and JCS performed data analysis. LC and LG performed IF. Histologic images were evaluated by LN, CW and DJ. Patient recruitment and care was performed by JG, MG, FI, SS, JS and KA. AC and SV generated and analyzed microCT data. Sample procurement was performed and supervised by JCK, CW, DJ and LN. The web portal was developed by SB and SG. CF, JG, KA, FI, BV, JH, AÖY, JGS, MMH, and NK provided critical interpretation, annotation, and comment on data and the manuscript. JR, HY, JCK and JCS drafted the manuscript, which was reviewed, edited, and approved by all authors.

## Supplemental Figures

**Figure S1. Tissue multiarrays (TMAs) and supplementary lineage annotation.**

A) Overview of hematoxylin/Eosin (HE) and Elastica van Gieson (EvG) stains of TMAs are analyzed in this study. 2 non-BOS / non-CTRL samples and 1 detached CTRL sample were excluded from downstream analysis yielding a total of s=108 samples.

B) Unity scaled average expression of canonical markers per sample across lineages within the snRNAseq dataset (10X Flex). Samples were grouped after disease and subject. Each subject was assigned to a unique color.

C) Unity scaled average expression of canonical markers per subject across lineages within the spatial transcriptomics dataset (10X Xenium). Subjects were grouped per disease. Each subject was assigned to a unique color.

D) Cross-platform validation of spatial cell-type annotations against the snRNAseq (Flex) reference, shown as a heatmap of Spearman correlations between every Xenium and Flex cell type computed over a shared feature space of the top 20 discriminating genes per cell type. Cell types are grouped by lineage on both axes, with the analysis restricted to structural cell types and genes measurable on both platforms.

Scale bars denote 500µm.

**Figure S2. Supplementary analysis of epithelial and mesenchymal lineage.**

A) Uniform Manifold Approximation and Projection (UMAP) representation of 10X Xenium data of the epithelial cell lineage, color coded by disease and lesion sub-class.

B) Bar plots show mean ± SD of cell-type proportions within the epithelial lineage. Variability across sub-classes was assessed by Kruskal-Walli’s test (*), with post hoc comparisons by Benjamini-Hochberg-adjusted Wilcoxon rank-sum test. Adjusted P < 0.05 was considered statistically significant (**Table S14-15**).

C) Unity scaled average expression of canonical epithelial markers per sample across epithelial cell types within the spatial transcriptomics dataset (10X Xenium). Samples were grouped after subject and disease. Each subject was assigned to a unique color.

D) Uniform Manifold Approximation and Projection (UMAP) representation of 10X Xenium data of the mesenchymal cell lineage, color coded by disease and sub-class.

E) Bar plots show mean ± SD of cell-type proportions within the mesenchymal lineage. Variability across sub-classes was assessed by Kruskal-Wallis test (*), with post hoc comparisons by Benjamini-Hochberg-adjusted Wilcoxon rank-sum test. Adjusted P < 0.05 was considered statistically significant (**Table S18-19**).

F) Unity scaled average expression of canonical mesenchymal markers per sample across mesenchymal cell types within the spatial transcriptomics dataset (10X Xenium). Samples were grouped after subject and disease. Each subject was assigned to a unique color.

**Figure S3. Array of epithelial cell types per sample.**

A) Spatial plot array shows of epithelial cells colored by cell type.

**Figure S4. Single channels of shown immune fluorescence images.**

A) - C) Single channel images of IF staining of SFTPC (green), CTSE (red), KRT17 (pink) and DAPI (blue). The merged channel images are displayed in Fig.2 N).

D) Single channel images of IF staining of PDPN (green), COL15A1 (red) and DAPI (blue). The merged channel images is displayed in Fig.4 K).

**Figure S5. Array of mesenchymal cell types per sample.**

A) Spatial plot array shows mesenchymal cells colored by cell type.

**Figure S6. Supplementary analysis of endothelial lineage.**

A) Uniform Manifold Approximation and Projection (UMAP) representation of 10X Xenium data of the endothelial cell lineage, color coded by disease and sub-class.

B) Bar plots show mean ± SD of cell-type proportions within the endothelial lineage. Variability across sub-classes was assessed by Kruskal-Wallis test (*), with post hoc comparisons by *Benjamini-Hochberg*-adjusted *Wilcoxon rank-sum* test. Adjusted P < 0.05 was considered statistically significant (**Table S23-24**).

C) Unity scaled average expression of canonical endothelial markers per sample across endothelial cell types within the spatial transcriptomics dataset (10X Xenium). Samples were grouped after subject and disease. Each subject was assigned to a unique color.

**Figure S7. Array of endothelial cell types per sample.**

A) Spatial plot array shows endothelial cells colored by cell type.

**Figure S8. Figure S6. Supplementary analysis of myeloid and lymphoid lineage.**

A) Uniform Manifold Approximation and Projection (UMAP) representation of 10X Xenium data of the myeloid cell lineage, color coded by disease and sub-class.

B) Bar plots show mean ± SD of cell-type proportions within the myeloid lineage. Variability across sub-classes was assessed by *Kruskal-Wallis* test (*), with post hoc comparisons by *Benjamini-Hochberg*-adjusted *Wilcoxon rank-sum* test. Adjusted P < 0.05 was considered statistically significant (**Table S25-26**).

C) Unity scaled average expression of canonical myeloid markers per sample across myeloid cell types within the spatial transcriptomics dataset (10X Xenium). Samples were grouped after subject and disease. Each subject was assigned to a unique color.

D) Uniform Manifold Approximation and Projection (UMAP) representation of 10X Xenium data of the lymphoid cell lineage, color coded by disease and sub-class.

E) Bar plots show mean ± SD of cell-type proportions within the lymphoid lineage. Variability across sub-classes was assessed by *Kruskal-Wallis* test (*), with post hoc comparisons by Benjamini-Hochberg-adjusted *Wilcoxon rank-sum* test. Adjusted P < 0.05 was considered statistically significant (**Table S27-28**).

F) Unity scaled average expression of canonical lymphoid markers per sample across lymphoid cell types within the spatial transcriptomics dataset (10X Xenium). Samples were grouped after subject and disease. Each subject was assigned to a unique color.

**Figure S9. Array of myeloid cell types per sample.**

A) Spatial plot array shows myeloid cells colored by cell type.

**Figure S10. Array of lymphoid cell types per sample.**

A) Spatial plot array shows lymphoid cells colored by cell type.

**Figure S11. Array of pan-lineage niches per sample.**

A) Spatial plot array shows all cells per samples colored by pan-lineage niches (N1-N7).

**Figure S12. LIANA+ spatially informed ligand-receptor (LR) analysis of the BOS lesion.**

A) Number of specific predicted ligand-receptor interactions per lesion sub-class, scored by the *LIANA+* rank aggregate consensus (specificity rank ≤ 0.05).

B) Mean bivariate Moran’s R of spatially informed LR interactions per section, compared between CTRL and BOS (Wilcoxon rank-sum test). Boxplots represent median and interquartile range, whiskers the 1.5xIQR.

C) Mean bivariate Moran’s R across sub-classes. Inter-group variability was assessed by *Kruskal-Wallis* test with *Benjamini-Hochberg-corrected* pairwise post hoc comparisons.

D) Heatmap of per-section ligand-receptor interaction strength (*LIANA+* spatially weighted local co-expression score), z-scored across sections per LR pair, with LR pairs clustered (left dendrogram)

E) Representative spatial plots of selected ligand-receptor pairs (*CXCL9*–*ACKR1*, *FN1*–*CD79A*, *CD80*–*CD28*) across *Inflamed*, *Obliterating* and *AFE* sections, colored by local interaction score on a scale shared across the displayed sections of a given pair.

F) -K) Chord diagrams of the top 10 edges per ligand-receptor pair. Rim sectors represent cell types and ribbons individual source-to-target interactions, with ribbon width scaling with interaction strength (−log10 magnitude rank), ribbon color denoting the sending cell type and arrowheads marking the receptor end.

Scale bars denote 500µm, unless stated otherwise.

## Supplemental Tables

**Table S1. Lineage marker genes per sample (snRNAseq).** Differentially expressed marker genes defining each structural lineage in the 10X Flex dataset, computed per sample by Wilcoxon rank-sum testing with *Bonferroni* correction.

**Table S2. Cell type marker genes per sample (snRNAseq).** Differentially expressed marker genes defining each structural cell type in the 10X Flex dataset, computed per sample by Wilcoxon rank-sum testing with *Bonferroni* correction.

**Table S3. Lineage composition across lesion sub-classes (Xenium).** Mean and standard deviation of relative lineage frequencies per lesion sub-class across all s=108 spatial transcriptomics samples.

**Table S4. Statistics of lineage composition across lesion sub-classes (Xenium).** *Kruskal-Wallis* tests and pairwise Wilcoxon rank-sum *post hoc* comparisons of lineage frequencies between lesion sub-classes, with *Benjamini-Hochberg* correction.

**Table S5. Lineage marker genes per subject (Xenium).** Marker genes defining each lineage in the spatial transcriptomics dataset, computed per subject by *Bonferroni*-adjusted Wilcoxon rank-sum testing.

**Table S6. Cell type marker genes per sample (Xenium).** Marker genes defining each annotated cell type in the spatial transcriptomics dataset, computed per sample by *Bonferroni*-adjusted Wilcoxon rank-sum testing.

**Table S7. Lineage frequencies per disease (snRNAseq).** Relative frequencies of structural lineages per subject in CTRL, BOS and RAS lungs.

**Table S8. Statistics of lineage frequencies per disease (snRNAseq).** *Kruskal-Wallis* tests and pairwise Wilcoxon rank-sum post hoc comparisons of lineage frequencies between CTRL, BOS and RAS, with *Benjamini-Hochberg* correction.

**Table S9. Epithelial cell type frequencies per disease (snRNAseq).** Relative frequencies of each epithelial cell type within the epithelial compartment per subject, stratified by disease.

**Table S10. Statistics of epithelial cell type frequencies per disease (snRNAseq).** *Kruskal-Wallis* tests and pairwise Wilcoxon rank-sum post hoc comparisons of epithelial cell type frequencies between CTRL, BOS and RAS, with *Benjamini-Hochberg* correction.

**Table S11. Pseudobulk differential gene expression across diseases (snRNAseq).** edgeR quasi-likelihood results for the contrasts BOS versus CTRL and BOS versus RAS, computed per cell type on subject-level pseudobulk profiles.

**Table S12. Cell type densities across lesion sub-classes (Xenium).** Mean and standard deviation of cell type densities per mm² of segmented tissue surface, stratified by lesion sub-class.

**Table S13. Statistics of cell type densities across lesion sub-classes (Xenium).** *Kruskal-Wallis* tests and pairwise Wilcoxon rank-sum post hoc comparisons of cell type densities between lesion sub-classes, with *Benjamini-Hochberg* correction.

**Table S14. Epithelial cell type composition across lesion sub-classes (Xenium).** Mean and standard deviation of relative epithelial cell type frequencies within the epithelial lineage per lesion sub-class.

**Table S15. Statistics of epithelial cell type composition across lesion sub-classes (Xenium).** *Kruskal-Wallis* tests and pairwise Wilcoxon rank-sum post hoc comparisons of epithelial cell type frequencies between lesion sub-classes, with *Benjamini-Hochberg* correction.

**Table S16. Mesenchymal cell type frequencies per disease (snRNAseq).** Relative frequencies of each mesenchymal cell type within the mesenchymal compartment per subject, stratified by disease.

**Table S17. Statistics of mesenchymal cell type frequencies per disease (snRNAseq).** *Kruskal-Wallis* tests and pairwise Wilcoxon rank-sum post hoc comparisons of mesenchymal cell type frequencies between CTRL, BOS and RAS, with *Benjamini-Hochberg* correction.

**Table S18. Mesenchymal cell type composition across lesion sub-classes (Xenium).** Mean and standard deviation of relative mesenchymal cell type frequencies within the mesenchymal lineage per lesion sub-class.

**Table S19. Statistics of mesenchymal cell type composition across lesion sub-classes (Xenium).** *Kruskal-Wallis* tests and pairwise Wilcoxon rank-sum post hoc comparisons of mesenchymal cell type frequencies between lesion sub-classes, with *Benjamini-Hochberg* correction.

**Table S20. Statistics of broncho-vascular niche surface proportions across lesion sub-classes (Xenium).** *Kruskal-Wallis* tests and pairwise Wilcoxon rank-sum *post hoc* comparisons of the relative surface proportions of the three niches constituting the broncho-vascular bundle, renormalized to 100% per sample and corrected by *Benjamini-Hochberg*.

**Table S21. Endothelial cell type frequencies per disease (snRNAseq).** Relative frequencies of each endothelial cell type within the endothelial compartment per subject, stratified by disease.

**Table S22. Statistics of endothelial cell type frequencies per disease (snRNAseq).** *Kruskal-Wallis* tests and pairwise Wilcoxon rank-sum post hoc comparisons of endothelial cell type frequencies between CTRL, BOS and RAS, with *Benjamini-Hochberg* correction.

**Table S23. Endothelial cell type composition across lesion sub-classes (Xenium).** Mean and standard deviation of relative endothelial cell type frequencies within the endothelial lineage per lesion sub-class.

**Table S24. Statistics of endothelial cell type composition across lesion sub-classes (Xenium).** *Kruskal-Wallis* tests and pairwise Wilcoxon rank-sum post hoc comparisons of endothelial cell type frequencies between lesion sub-classes, with *Benjamini-Hochberg* correction.

**Table S25. Myeloid cell type composition across lesion sub-classes (Xenium).** Mean and standard deviation of relative myeloid cell type frequencies within the myeloid lineage per lesion sub-class.

**Table S26. Statistics of myeloid cell type composition across lesion sub-classes (Xenium).** *Kruskal-Wallis* tests and pairwise Wilcoxon rank-sum post hoc comparisons of myeloid cell type frequencies between lesion sub-classes, with *Benjamini-Hochberg* correction.

**Table S27. Lymphoid cell type composition across lesion sub-classes (Xenium).** Mean and standard deviation of relative lymphoid cell type frequencies within the lymphoid lineage per lesion sub-class.

**Table S28. Statistics of lymphoid cell type composition across lesion sub-classes (Xenium).** *Kruskal-Wallis* tests and pairwise Wilcoxon rank-sum post hoc comparisons of lymphoid cell type frequencies between lesion sub-classes, with *Benjamini-Hochberg* correction.

**Table S29. Quality control metrics of the snRNAseq dataset.** Per-sample sequencing and quality control metrics of the 10X Flex libraries.

**Table S30. Custom Xenium add-on panel gene list.** The 100 custom target genes added to the Xenium Human Lung Panel v1, selected from the snRNAseq data by cell type discriminative power.

**Table S31. Antibodies used for multiplex immunofluorescence.** Primary and secondary antibodies used for protein-level validation.

## References

1. Verleden, G. M. et al. Chronic lung allograft dysfunction: Definition, diagnostic criteria, and approaches to treatment—A consensus report from the Pulmonary Council of the ISHLT. The Journal of Heart and Lung Transplantation 38, 493–503 (2019).

2. Estenne, M. & Hertz, M. I. Bronchiolitis Obliterans after Human Lung Transplantation. American Journal of Respiratory and Critical Care Medicine 166, 440–444 (2002).

3. Verleden, S. E. et al. When tissue is the issue: A histological review of chronic lung allograft dysfunction. American Journal of Transplantation 20, 2644–2651 (2020).

4. Verleden, S. E., Vos, R., Vanaudenaerde, B. M. & Verleden, G. M. Chronic lung allograft dysfunction phenotypes and treatment. Journal of Thoracic Disease 9, 2650–2659 (2017).

5. Mellors, P. W., et al. Shared roles of immune and stromal cells in the pathogenesis of human bronchiolitis obliterans syndrome. JCI Insight 10, e176596 (2025).

6. Bos, S., Milross, L., Filby, A. J., Vos, R. & Fisher, A. J. Immune processes in the pathogenesis of chronic lung allograft dysfunction: identifying the missing pieces of the puzzle. European Respiratory Review 31, 220060 (2022).

7. Sato, M. Bronchiolitis obliterans syndrome and restrictive allograft syndrome after lung transplantation: why are there two distinct forms of chronic lung allograft dysfunction? Annals of Translational Medicine 8, 418 (2020).

8. Khatri, A. et al. JAK-STAT activation contributes to cytotoxic T cell–mediated basal cell death in human chronic lung allograft dysfunction. JCI Insight 8, e167082 (2023).

9. Yan, Y., et al. Single-cell dissection of chronic lung allograft dysfunction reveals convergent and distinct fibrotic mechanisms. JCI Insight 10, (2025).

10. Ruwisch, J. et al. The pleuroparenchymal fibroelastosis atlas reveals aberrant cell states and their zonation as an alternate roadmap to lung fibrosis. Science Advances 12, eaeb5967 (2026).

11. Braubach, P. et al. Pulmonary fibroelastotic remodelling revisited. Cells 10, (2021).

12. Leiber, L. M. et al. Aberrant and Ectopic Cell Populations in Restrictive Allograft Syndrome after Lung Transplantation. Eur Respir J 2500537 (2026) doi:10.1183/13993003.00537-2025.

13. Guo, M. et al. Guided construction of single cell reference for human and mouse lung. Nat Commun 14, 4566 (2023).

14. Sikkema, L. et al. An integrated cell atlas of the lung in health and disease. Nature Medicine 2023 29:6 29, 1563–1577 (2023).

15. Adams, T. S. et al. Single-cell RNA-seq reveals ectopic and aberrant lung-resident cell populations in idiopathic pulmonary fibrosis. Science Advances 6, eaba1983 (2020).

16. Schupp, J. C. et al. Integrated Single-Cell Atlas of Endothelial Cells of the Human Lung. Circulation 144, 286–302 (2021).

17. Adams, T. S. et al. Alveolar epithelial cell plasticity and injury memory in human pulmonary fibrosis. bioRxiv 2025.06.10.658504 (2025) doi:10.1101/2025.06.10.658504.

18. Varrone, M., Tavernari, D., Santamaria-Martínez, A., Walsh, L. A. & Ciriello, G. CellCharter reveals spatial cell niches associated with tissue remodeling and cell plasticity. Nat Genet 56, 74–84 (2024).

19. Tsukui, T. & Sheppard, D. Stromal heterogeneity in the adult lung delineated by single-cell genomics. American Journal of Physiology-Cell Physiology 328, C1964–C1972 (2025).

20. Tsukui, T., Wolters, P. J. & Sheppard, D. Alveolar fibroblast lineage orchestrates lung inflammation and fibrosis. Nature 631, 627–634 (2024).

21. Madissoon, E. et al. A spatially resolved atlas of the human lung characterizes a gland-associated immune niche. Nat Genet 55, 66–77 (2023).

22. Kadur Lakshminarasimha Murthy, P., et al. Human distal lung maps and lineage hierarchies reveal a bipotent progenitor. Nature 2022 604:7904 604, 111–119 (2022).

23. Fridman, W. H. et al. Tertiary lymphoid structures and B cells: An intratumoral immunity cycle. Immunity 56, 2254–2269 (2023).

24. Zhao, L. et al. Tertiary lymphoid structures in diseases: immune mechanisms and therapeutic advances. Signal Transduct Target Ther 9, 225 (2024).

25. Schumacher, T. N. & Thommen, D. S. Tertiary lymphoid structures in cancer. Science 375, eabf9419 (2022).

26. Fleige, H. et al. IL-17–induced CXCL12 recruits B cells and induces follicle formation in BALT in the absence of differentiated FDCs. J Exp Med 211, 643–651 (2014).

27. Pandey, S. et al. IL-4/CXCL12 loop is a key regulator of lymphoid stroma function in follicular lymphoma. Blood 129, 2507–2518 (2017).

28. Habermann, A. C. et al. Single-cell RNA sequencing reveals profibrotic roles of distinct epithelial and mesenchymal lineages in pulmonary fibrosis. Science Advances 6, (2020).

29. Vannan, A. et al. Spatial transcriptomics identifies molecular niche dysregulation associated with distal lung remodeling in pulmonary fibrosis. Nat Genet https://doi.org/10.1038/s41588-025-02080-x (2025) doi:10.1038/s41588-025-02080-x.

30. Mayr, C. H. et al. Spatial transcriptomic characterization of pathologic niches in IPF. Science Advances 10, eadl5473 (2024).

31. Schupp, J. C. et al. Alveolar Vascular Remodeling in Nonspecific Interstitial Pneumonia: Replacement of Normal Lung Capillaries with COL15A1 Positive Endothelial Cells. https://doi.org/10.1164/rccm.202303-0544LE https://doi.org/10.1164/RCCM.202303-0544LE (2023) doi:10.1164/RCCM.202303-0544LE.

32. Gottlieb, J. et al. Disease progression in patients with the restrictive and mixed phenotype of Chronic Lung Allograft dysfunction—A retrospective analysis in five European centers to assess the feasibility of a therapeutic trial. PLOS ONE 16, e0260881 (2021).

33. Joulia, R. et al. A single-cell spatial chart of the airway wall reveals proinflammatory cellular ecosystems and their interactions in health and asthma. Nat Immunol 26, 920–933 (2025).

34. Silva, M. V. et al. CD28 Family and Chronic Rejection: ‘To Belatacept…and Beyond!’ Journal of Transplantation 2012, 203780 (2012).

35. Mathews, D. V. et al. Belatacept-Resistant Rejection Is Associated With CD28+ Memory CD8 T Cells. American Journal of Transplantation 17, 2285–2299 (2017).

36. Thiriot, A. et al. Differential DARC/ACKR1 expression distinguishes venular from non-venular endothelial cells in murine tissues. BMC Biology 15, 45 (2017).

37. Guo, X. et al. Endothelial ACKR1 is induced by neutrophil contact and down-regulated by secretion in extracellular vesicles. Frontiers in Immunology 14, 1181016 (2023).

38. Butler, A., Hoffman, P., Smibert, P., Papalexi, E. & Satija, R. Integrating single-cell transcriptomic data across different conditions, technologies, and species. Nat Biotechnol 36, 411–420 (2018).

39. Gaddis, N. et al. LungMAP Portal Ecosystem: Systems-level Exploration of the Lung. Am J Respir Cell Mol Biol 70, 129–139 (2024).

40. Marco Salas, S., et al. Optimizing Xenium In Situ data utility by quality assessment and best-practice analysis workflows. Nat Methods 22, 813–823 (2025).

41. Quanjer, P. H. et al. MULTI-ETHNIC REFERENCE VALUES FOR SPIROMETRY FOR THE 3–95 YEAR AGE RANGE: THE GLOBAL LUNG FUNCTION 2012 EQUATIONS: Report of the Global Lung Function Initiative (GLI), ERS Task Force to establish improved Lung Function Reference Values. The European respiratory journal 40, 1324 (2012).

