## Supplementary_FiguresS1_to_S12 for "Single-cell and spatial transcriptomics resolve airway obliteration in bronchiolitis obliterans syndrome"

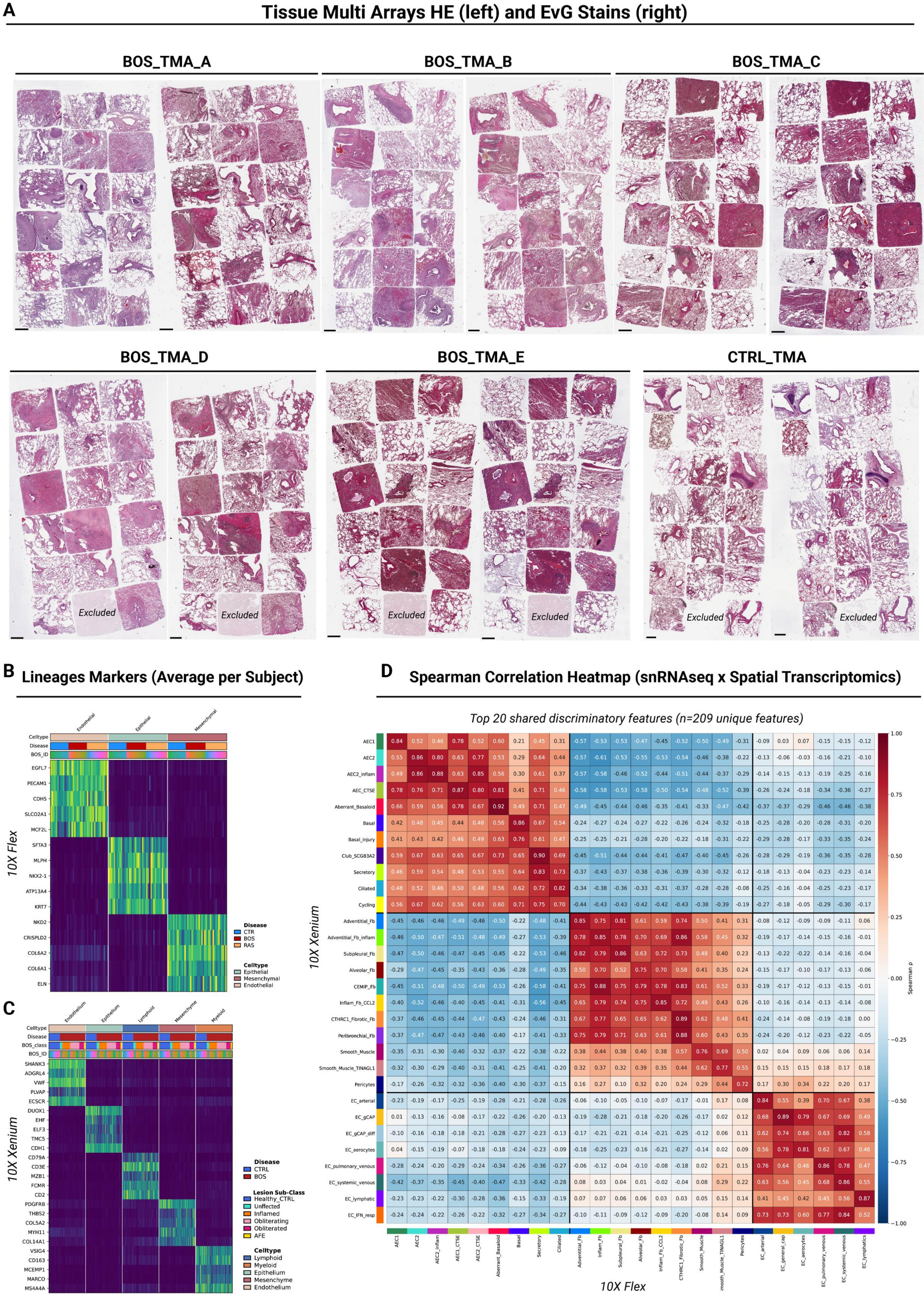

A
Tissue Multi Arrays HE (left) and EvG Stains (right)
BOS_TMA_A
BOS_TMA_B
BOS_TMA_C
BOS_TMA_D
BOS_TMA_E
CTRL_TMA
Excluded
B
Lineages Markers (Average per Subject)
10X Flex
Celltype
Disease
BOS_ID
Endothelial
Epithelial
Mesenchymal
EGFL7
PECAM1
CDH5
SLCO2A1
MCF2L
SFTA3
MLPH
NKX2-1
ATP13A4
KRT7
NKD2
CRISPLD2
COL6A2
COL6A1
ELN
Disease
CTR
BOS
RAS
Celltype
Epithelial
Mesenchymal
Endothelial
C
10X Xenium
Celltype
Disease
BOS_class
BOS_ID
Endothelium
Epithelium
Lymphoid
Mesenchyme
Myeloid
SHANK3
ADGRL4
VWF
PLVAP
ECSCR
DUOX1
EHF
ELF3
TMC5
CDH1
CD79A
CD3E
MZB1
FCMR
CD2
PDGFRB
THBS2
COL5A2
MYH11
COL14A1
VSIG4
CD163
MCEMP1
MARCO
MS4A4A
Disease
CTRL
BOS
Lesion Sub-Class
Healthy_CTRL
Unaffected
Inflamed
Obliterating
Obliterated
AFE
Celltype
Lymphoid
Myeloid
Epithelium
Mesenchyme
Endothelium
D
Spearman Correlation Heatmap (snRNAseq x Spatial Transcriptomics)
Top 20 shared discriminatory features (n=209 unique features)
10X Xenium
10X Flex
Spearman ρ

### Epithelial Lineage

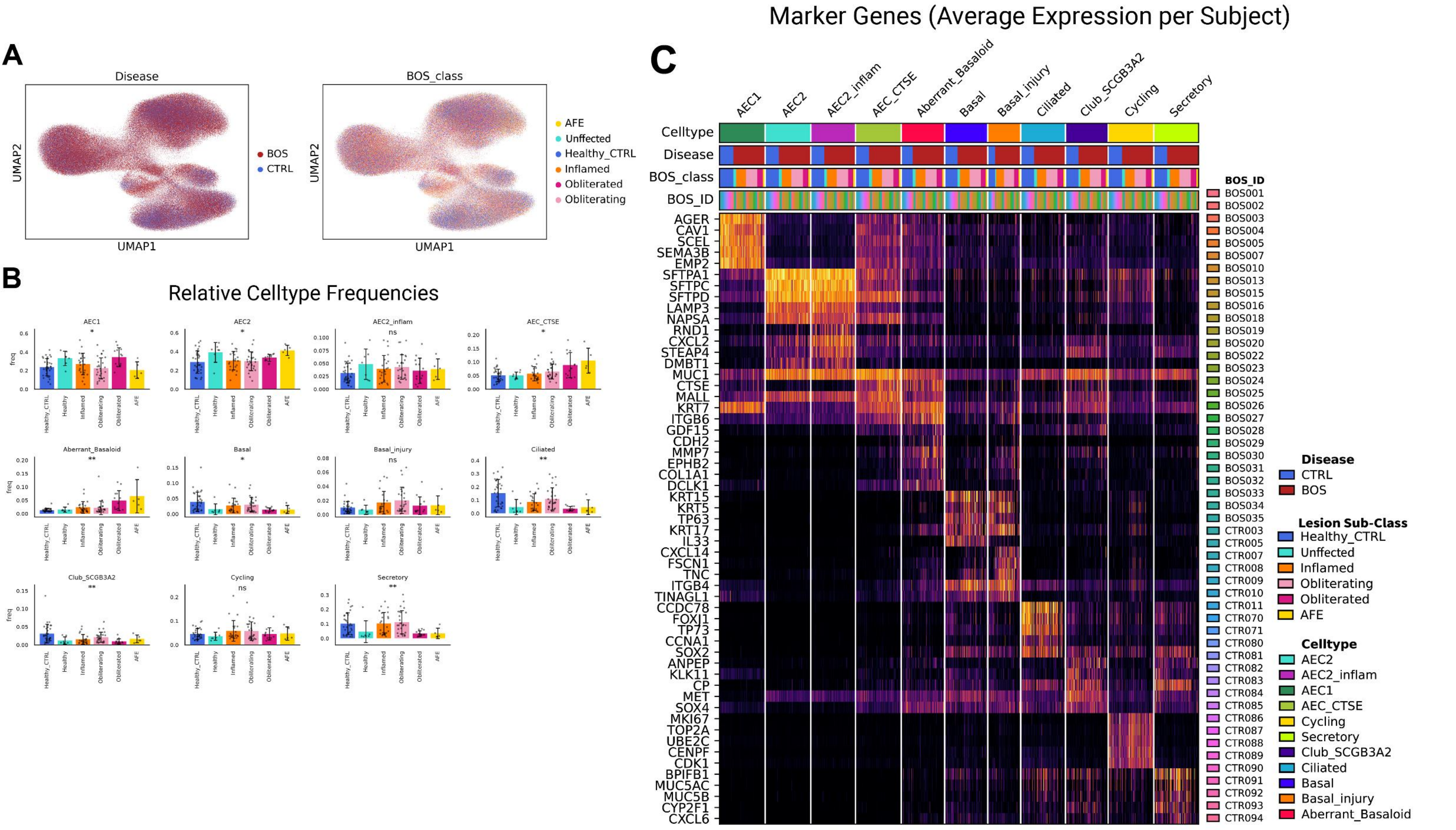


### Mesenchymal Lineage

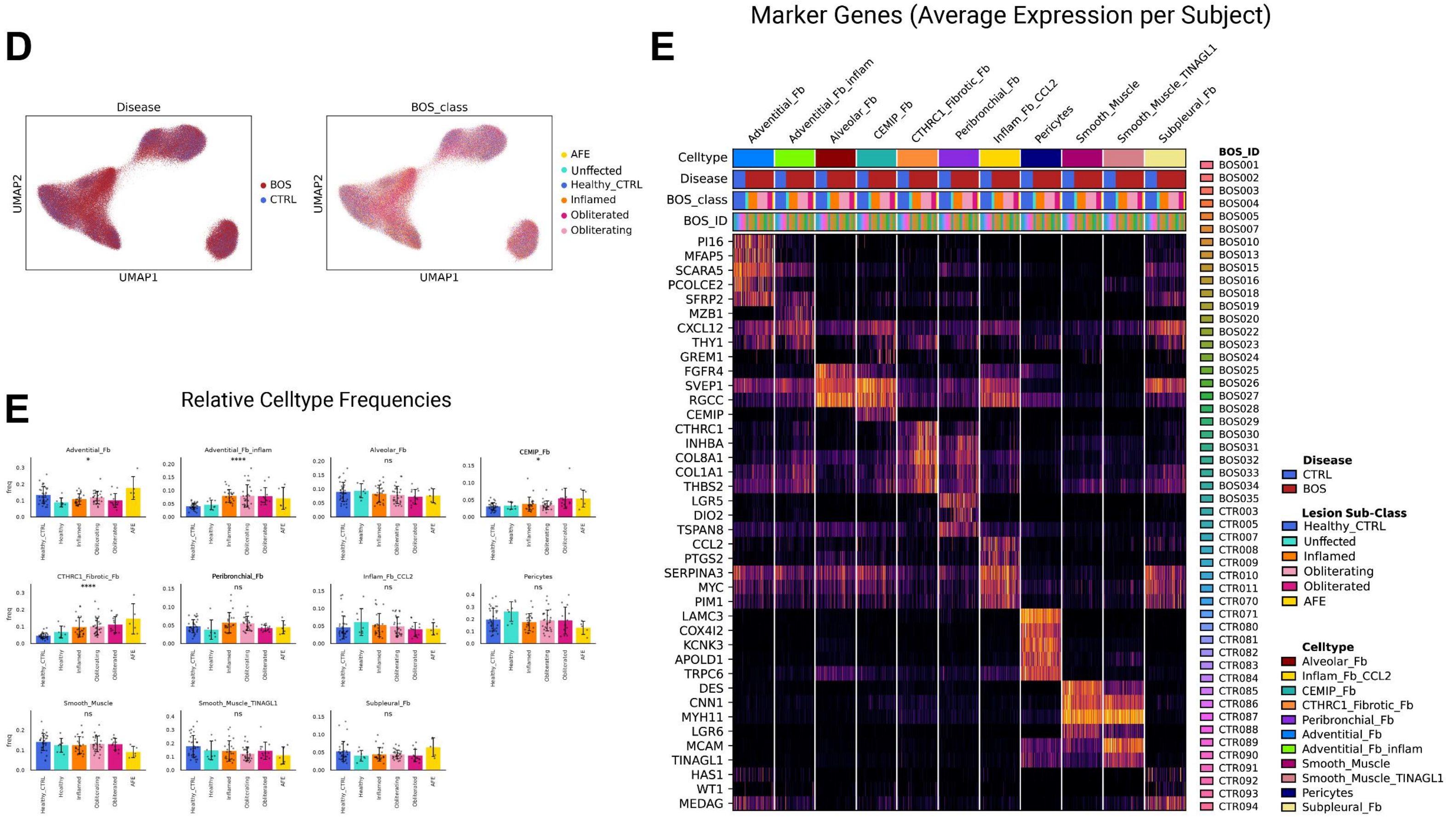

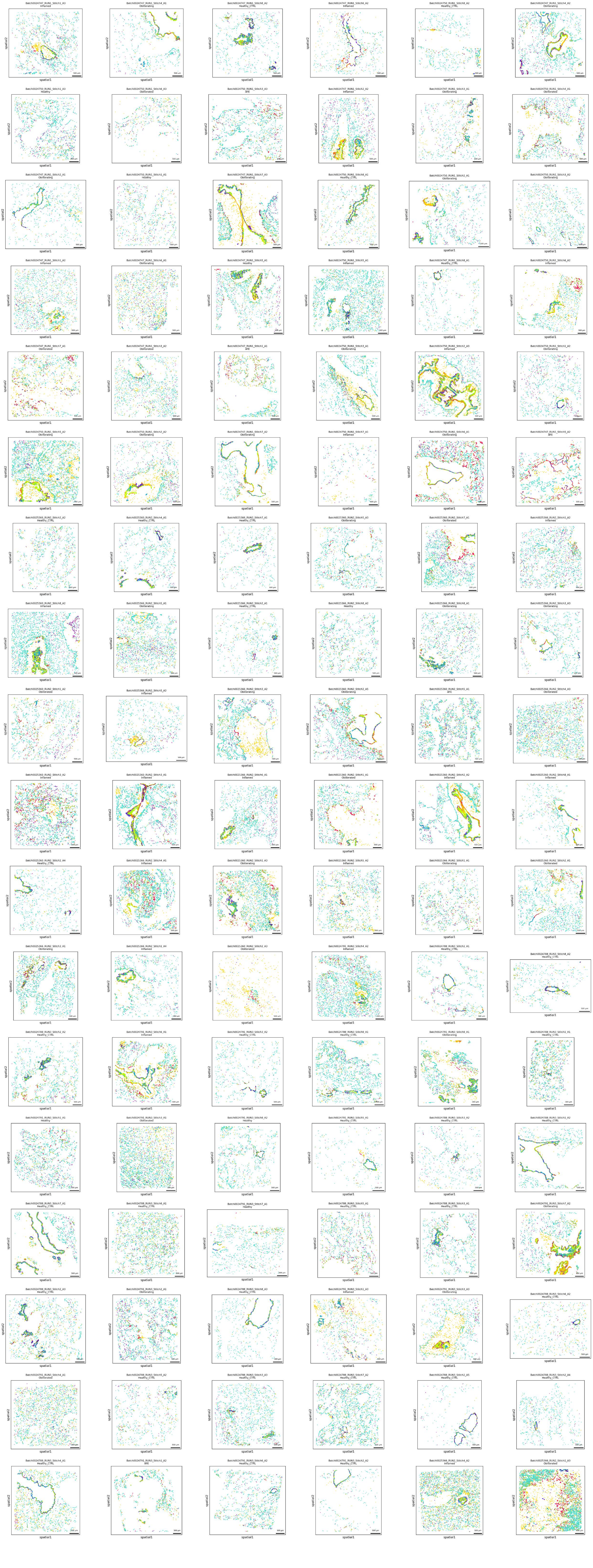
spatial1
spatial2
500 µm
AEC2
AEC2_inflam
AEC_CTSE
Cycling
Secretory
Club_SCGB3A2
Ciliated
Basal
Basal_injury
Aberrant_Basaloid

#### Protein Level (Single Channels)

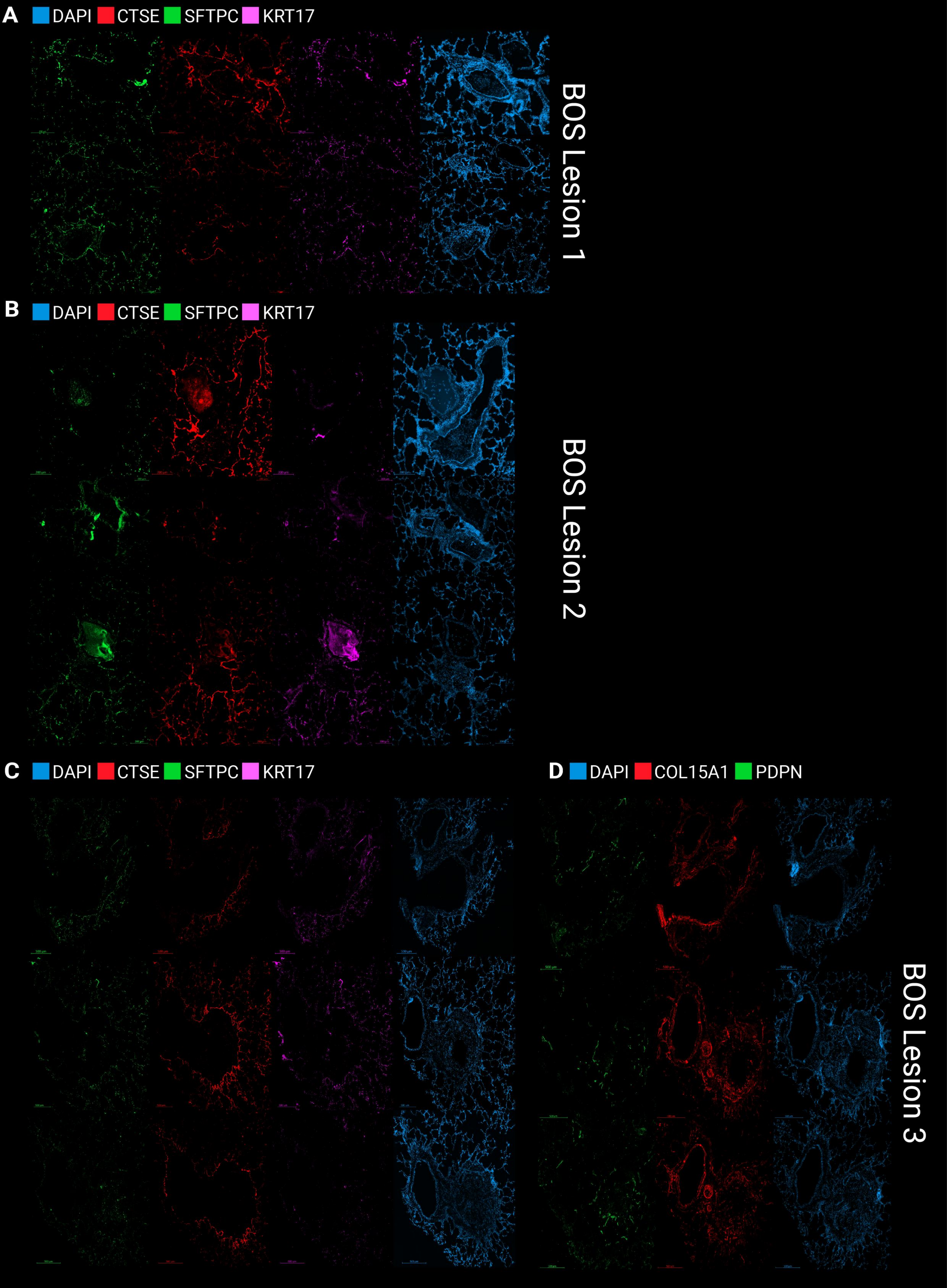

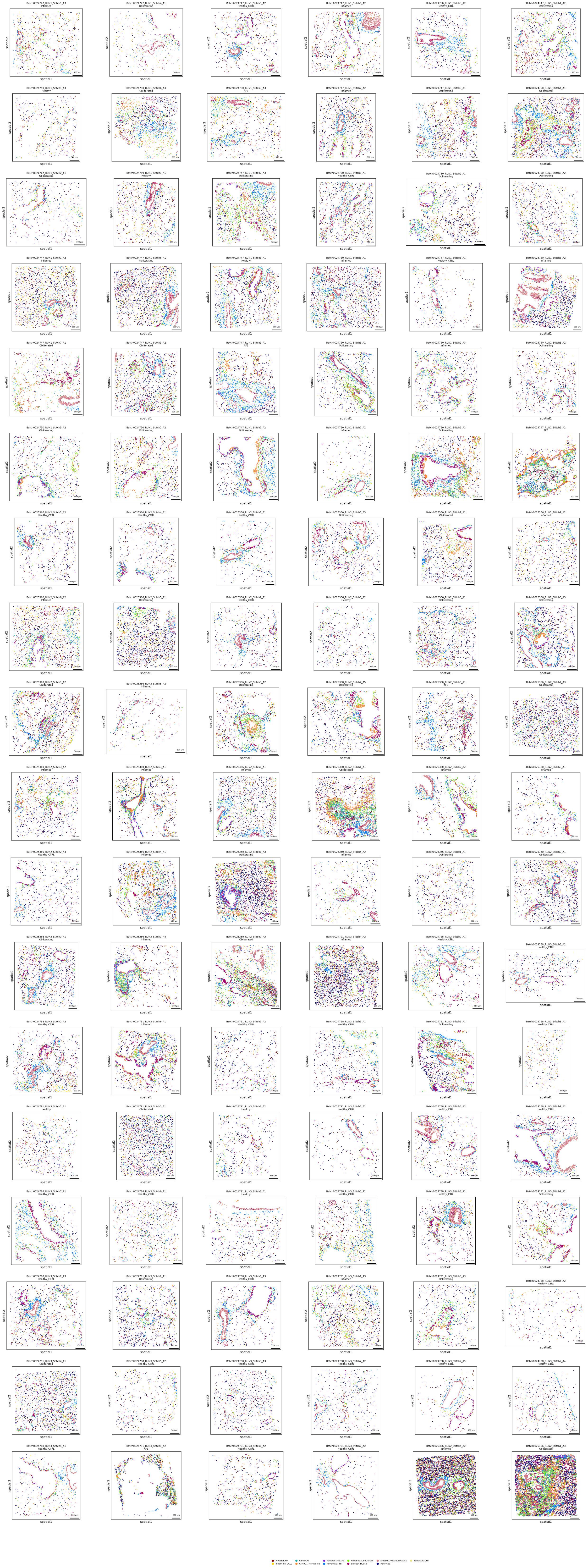
spatial1
spatial2
Alveolar_Fb
Inflam_Fb_CCL2
CEMIP_Fb
CTHRC1_Fibrotic_Fb
Peribronchial_Fb
Adventitial_Fb
Adventitial_Fb_inflam
Smooth_Muscle
Smooth_Muscle_TINAGL1
Pericytes
Subpleural_Fb

### Endothelial Lineage

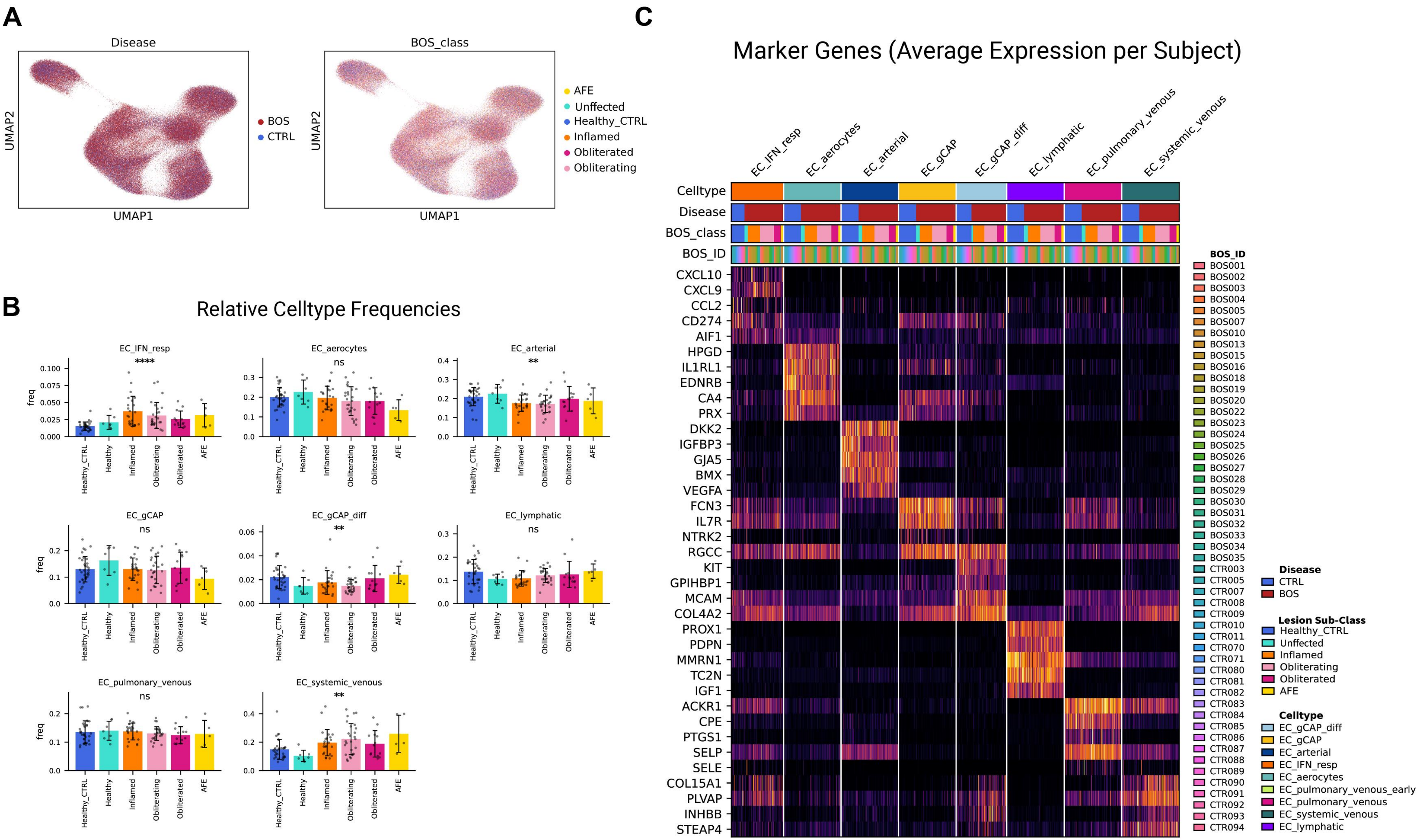

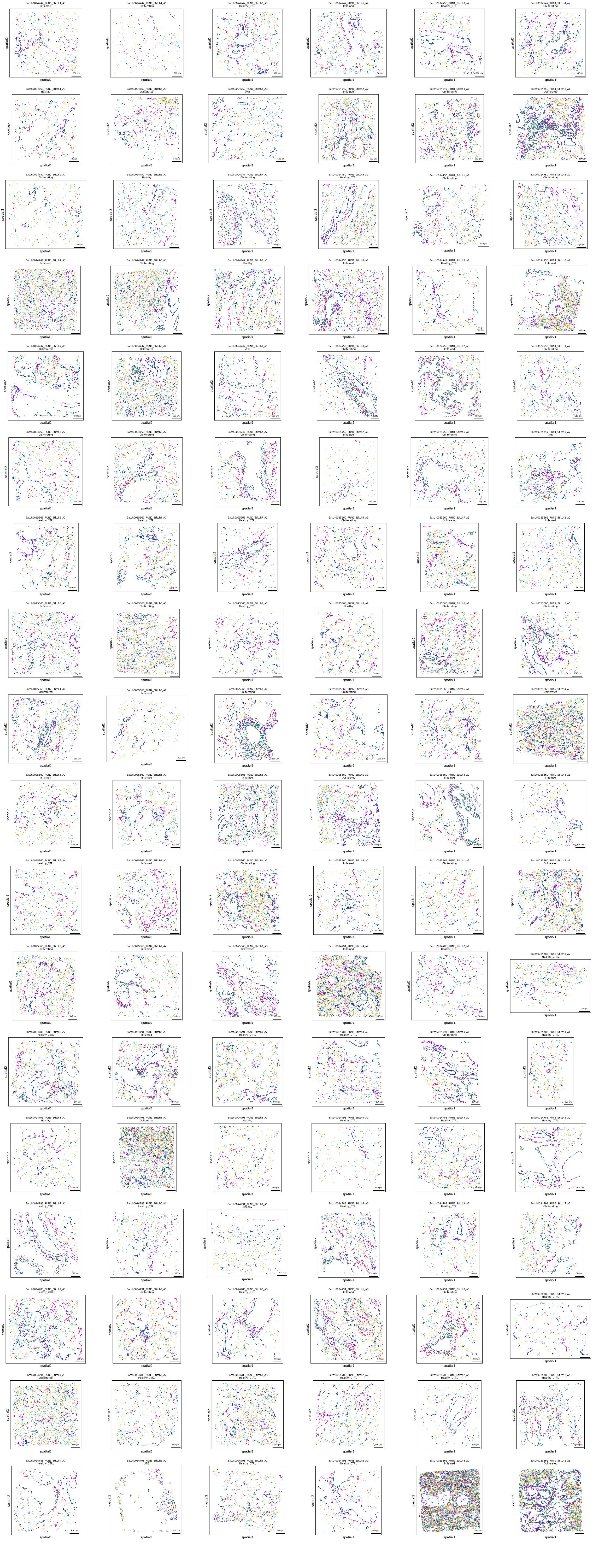
spatial1
spatial2
500 µm
EC_gCAP
EC_gCAP_diff
EC_arterial
EC_aerocytes
EC_pulmonary_venous
EC_systemic_venous
EC_lymphatic
EC_IFN_resp

### Myeloid Lineage

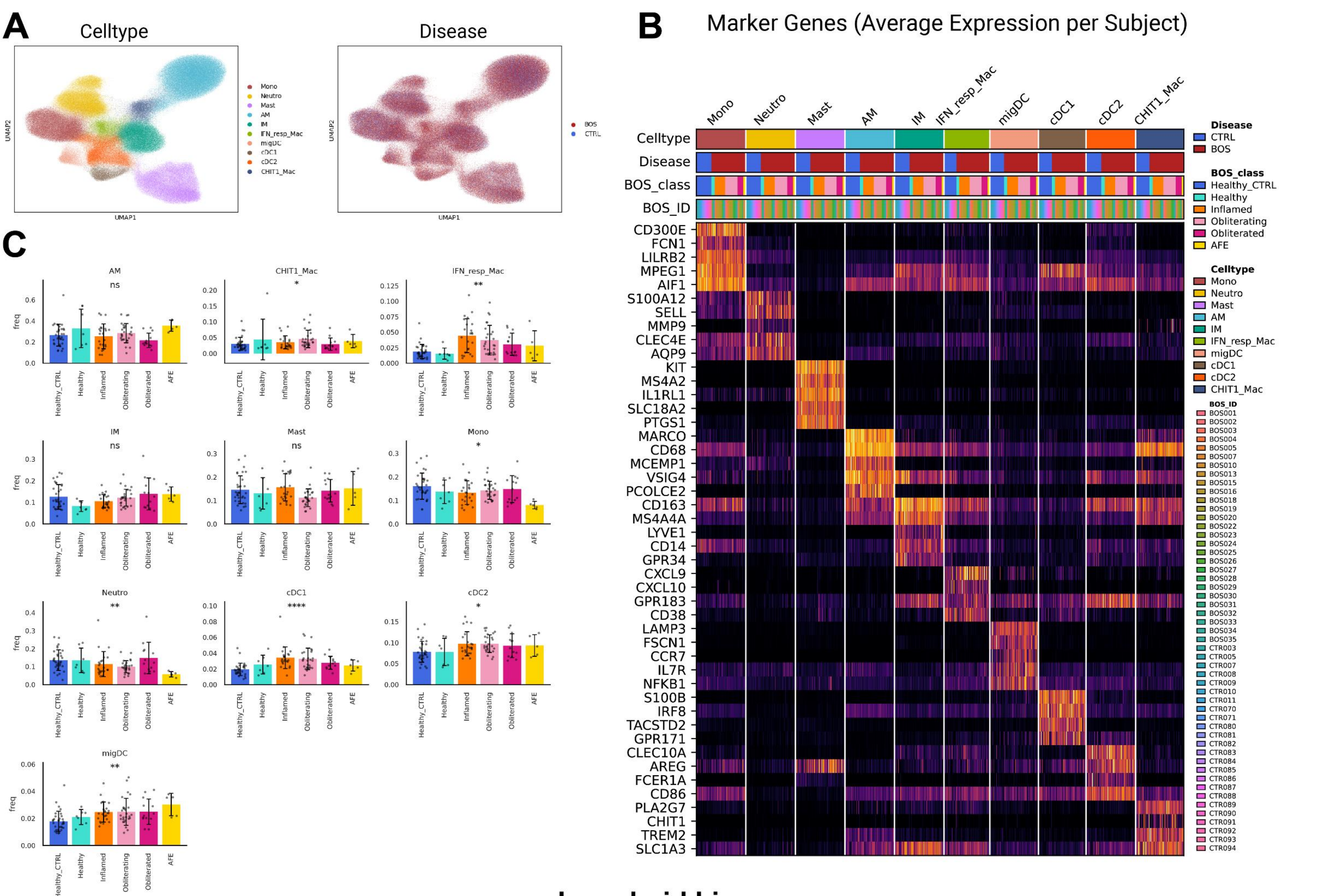


### Lymphoid Lineage

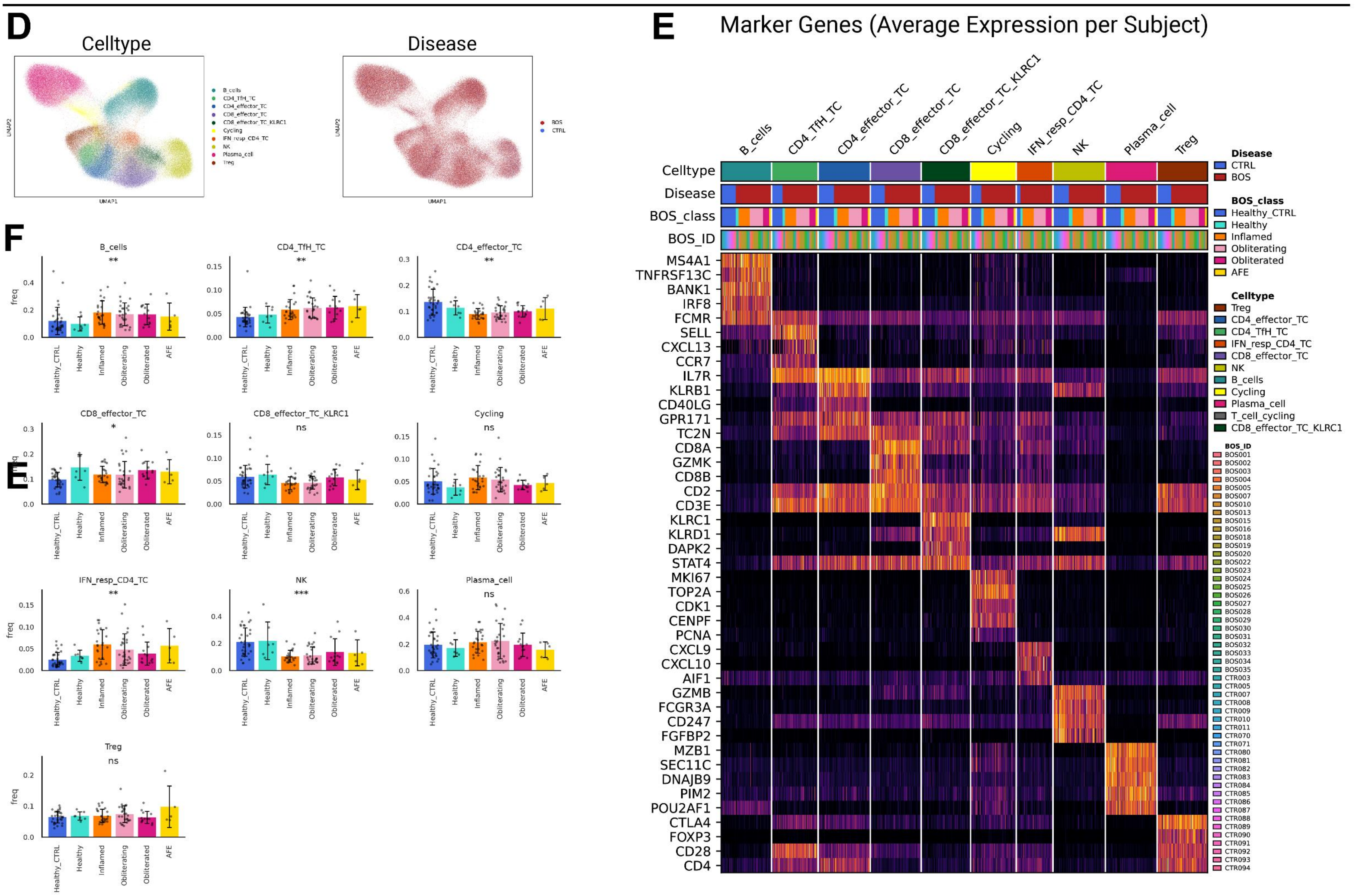

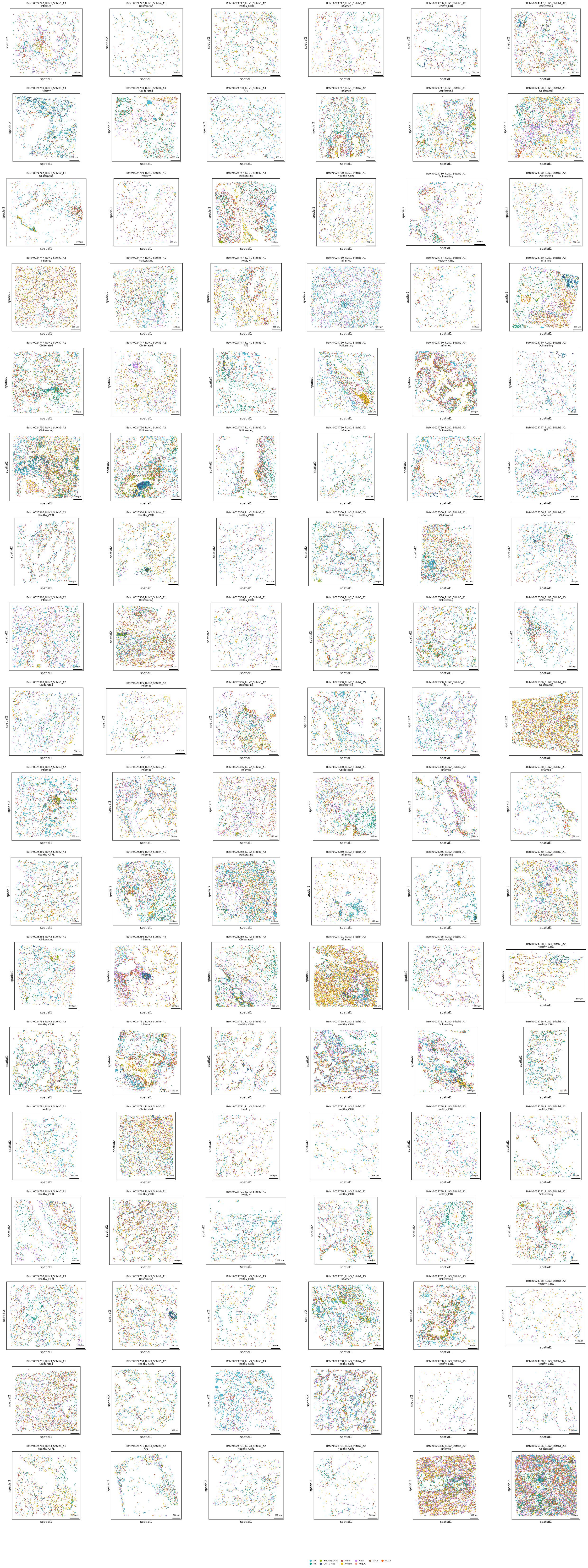
AM
IM
IFN_resp_Mac
CHIT1_Mac
Mono
Neutro
Mast
migDC
cDC1
cDC2

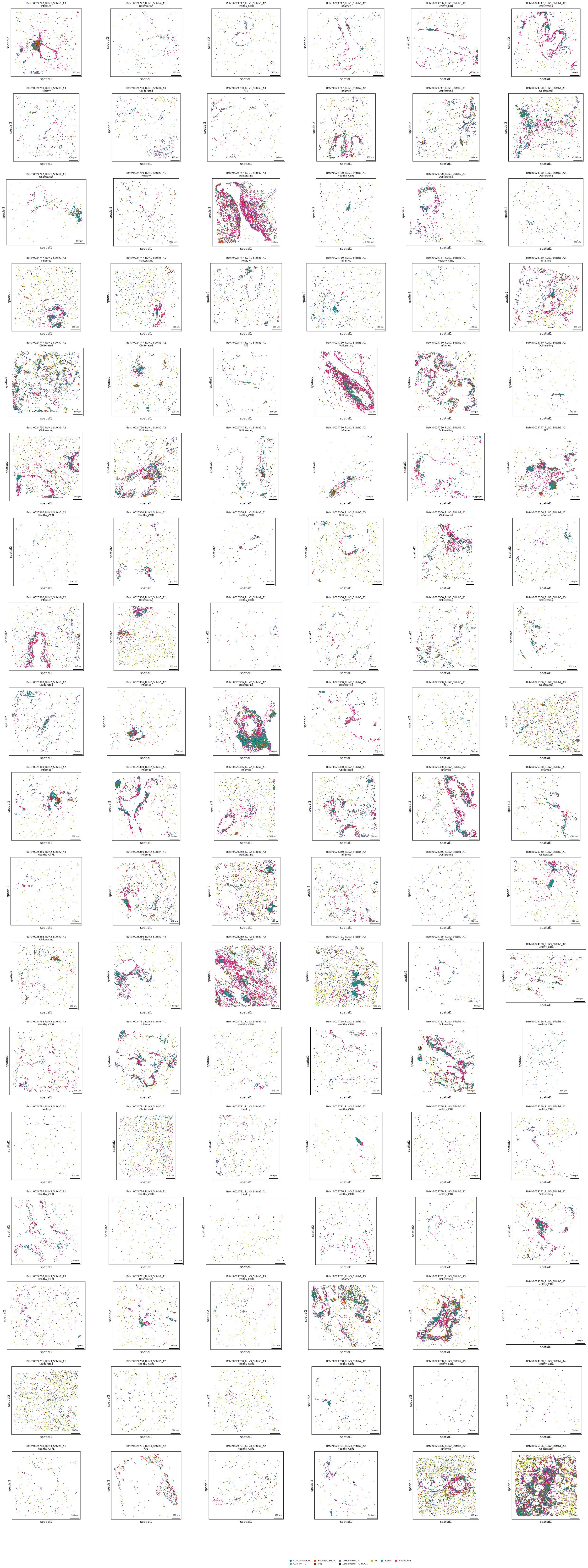
spatial1
spatial2
500 µm
CD4_effector_TC
CD4_TfH_TC
IFN_resp_CD4_TC
Treg
CD8_effector_TC
CD8_effector_TC_KLRC1
NK
B_cells
Plasma_cell

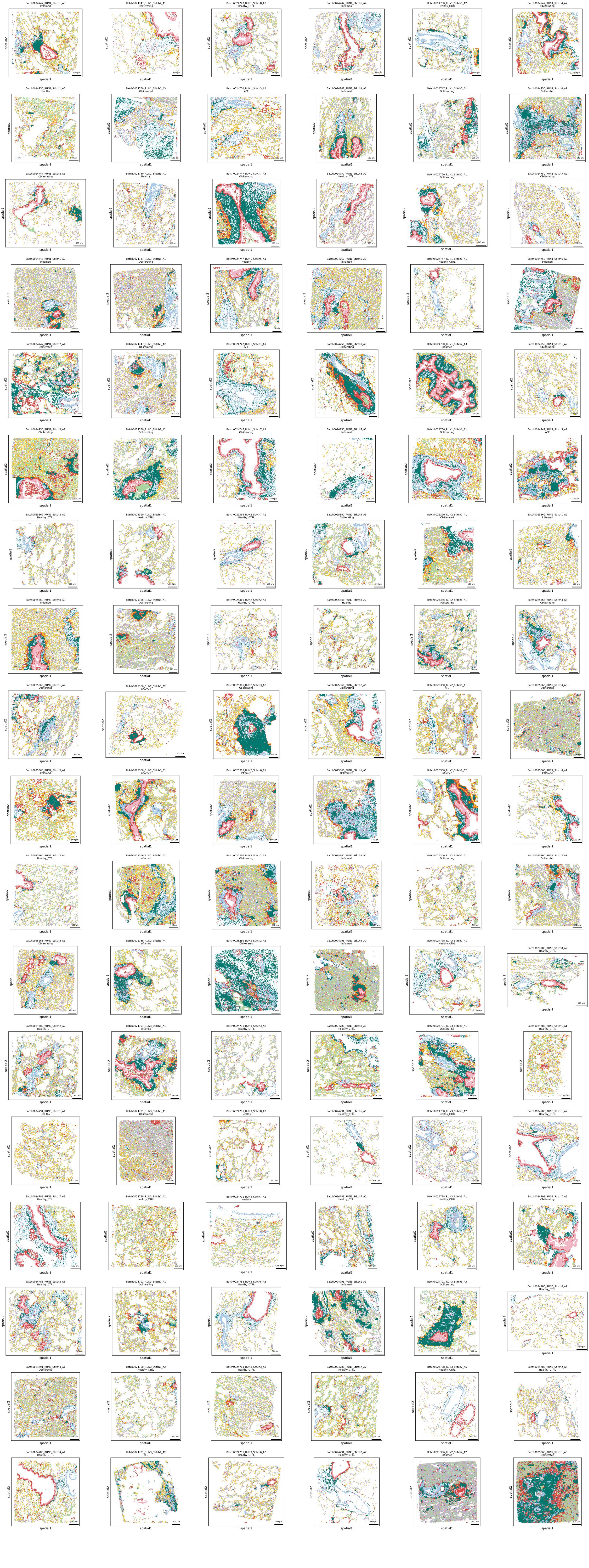
spatial1
spatial2
500 µm
Niche N1
Niche N2
Niche N3
Niche N4
Niche N5
Niche N6
Niche N7

### LIANA+ Spatially Informed Ligand-Receptor Analysis

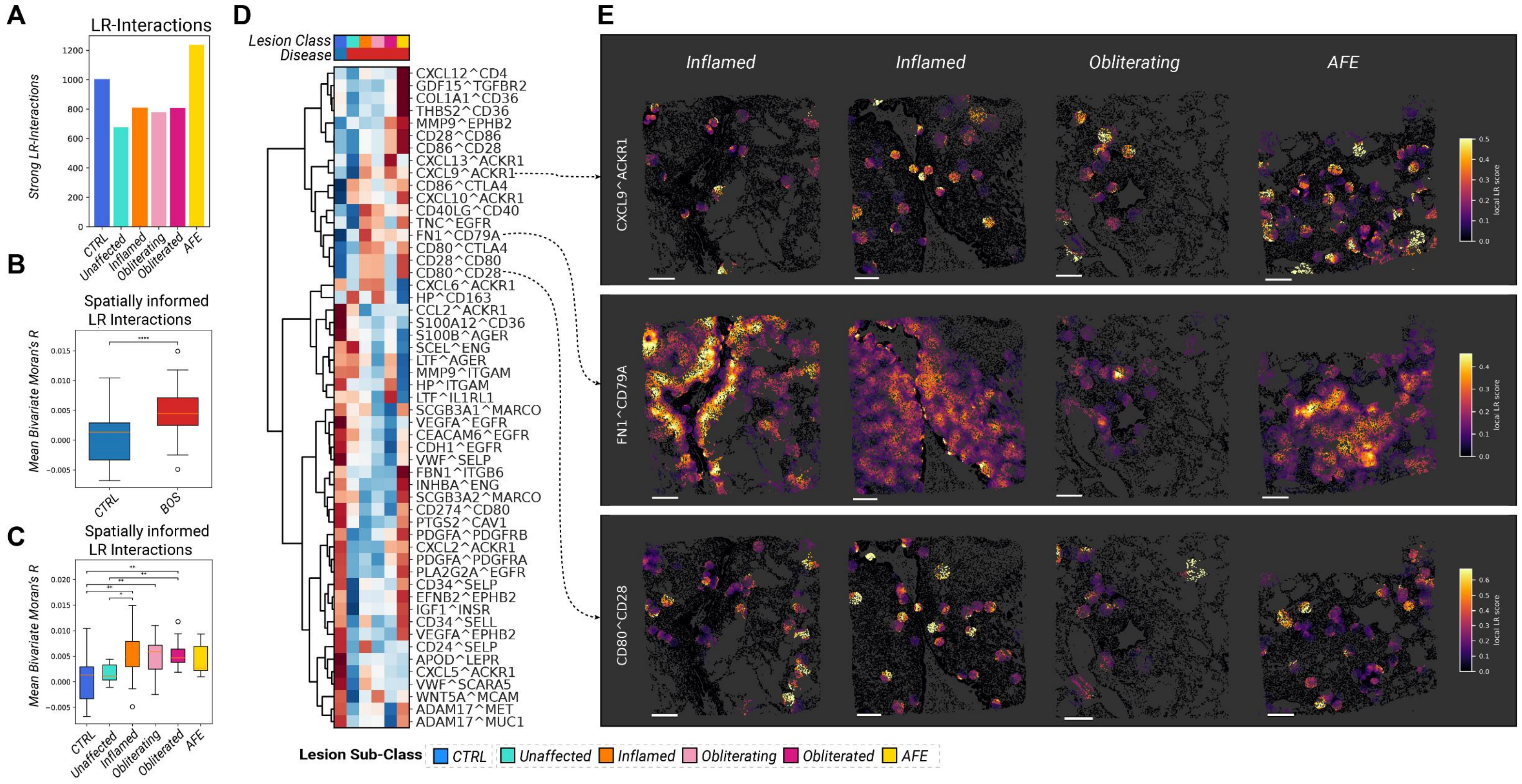


### LIANA+ Ligand Receptor Analysis (Top 10 Eges per LR-Pair)

Edge Tickness = LIANA+ Magnitude Rank

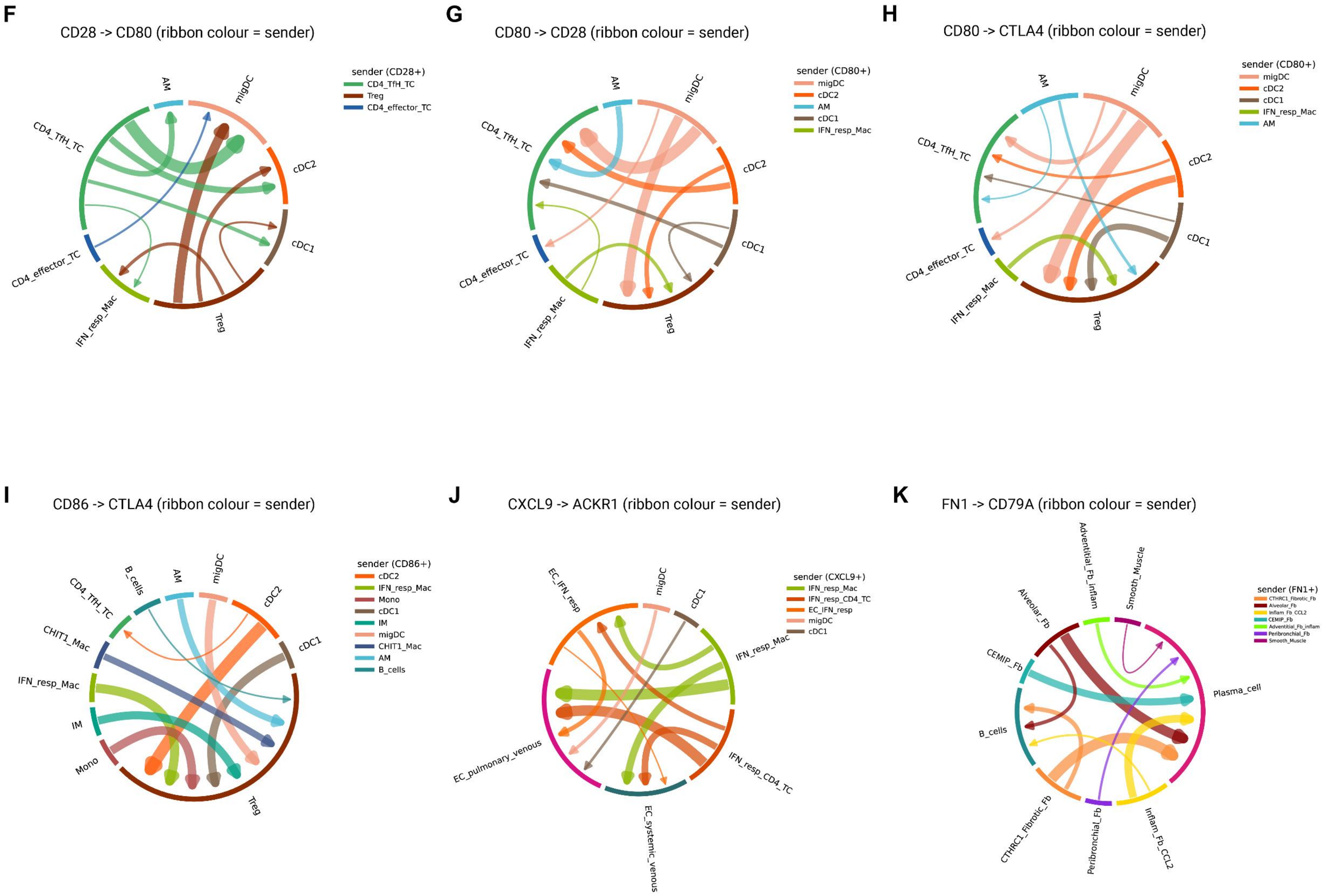
